# Dynamic flexibility in striatal-cortical circuits supports reinforcement learning

**DOI:** 10.1101/094383

**Authors:** Raphael T. Gerraty, Juliet Y. Davidow, Karin Foerde, Adriana Galvan, Danielle S. Bassett, Daphna Shohamy

## Abstract

Complex learned behaviors must involve the integrated action of distributed brain circuits. While the contributions of individual regions to learning have been extensively investigated, understanding how distributed brain networks orchestrate their activity over the course of learning remains elusive. To address this gap, we used fMRI combined with tools from dynamic network neuroscience to obtain time-resolved descriptions of network coordination during reinforcement learning. We found that learning to associate visual cues with reward involves dynamic changes in network coupling between the striatum and distributed brain regions, including visual, orbitofrontal, and ventromedial prefrontal cortex. Moreover, we found that flexibility in striatal network dynamics correlates with participants’ learning rate and inverse temperature, two parameters derived from reinforcement learning models. Finally, we found that not all forms of learning relate to this circuit: episodic memory, measured in the same participants at the same time, was related to dynamic connectivity in distinct brain networks. These results suggest that dynamic changes in striatal-centered networks provide a mechanism for information integration during reinforcement learning.

**Significance Statement:** Learning from the outcomes of actions–referred to as *reinforcement learning*–is an essential part of life. The roles of individual brain regions in reinforcement learning have been well characterized in terms of the updating of values for actions or sensory stimuli. Missing from this account, however, is a description of the manner in which different brain areas interact during learning to integrate sensory and value information. Here we characterize flexible striatal-cortical network dynamics that relate to reinforcement learning behavior.

## Introduction

Learning from reinforcement is central to adaptive behavior and requires continuous and dynamic integration of sensory, motor, and reward information over time. Major progress has been made in understanding how individual brain regions support reinforcement learning. However, remarkably little is known about how these brain regions interact during learning, how their interactions change over time, and how these dynamic circuit-level changes relate to successful learning.

In a typical reinforcement learning task, participants use reinforcement over hundreds of trials to associate cues or actions with their most probable outcome (e.g. (1–4)). Computationally, this is captured by so-called “model-free” reinforcement learning algorithms, a class of models that provide a quantitative and mechanistic framework for describing behavior (1, 5, 6). These models have also been successful in accounting for neuronal signals underlying learning behavior (3, 7, 8), demonstrating a role for the striatum and its dopaminergic inputs in updating reward predictions. However, to support reinforcement learning, the striatum must also integrate visual, motor, and reinforcement information over time. Such a process is likely to involve dynamic coordination across a number of different circuits interconnected with the striatum.

The idea that the striatum serves an integrative role in learning and cognition is not new (9–15). Anatomically, the striatum is well positioned for integration: it receives extensive input from many regions of cortex and projects back, through thalamus, to motor cortex (15–18). However, while the idea that the striatum serves such a role is anatomically and theoretically appealing, it has been very difficult to test this possibility empirically. Thus, whether the striatum interacts with other sensory, motor and cognitive regions during learning, and how these network-level interactions reconfigure over the course of learning, remains unknown.

Understanding the process of integration at the network level has been hampered by a lack of tools capable of quantifying the reconfiguration of those circuits in a data-driven fashion as humans adapt their behavior. Until now, most studies of large-scale brain connectivity have focused on static descriptions of networks (19–22), limiting their ability to link networks to cognitive processes (23). Yet, there is increasing recognition of the importance of a more dynamic perspective on circuit configuration. Even seemingly stable networks undergo temporal changes (24–26), implying that static descriptions fail to capture transient patterns of co-activation that may be essential for complex behavior.

Here we aimed to address this gap. We take advantage of recent advances in a dynamic formulation of graph theory and its application to neuroimaging data, an emerging field known as dynamic network neuroscience (23, 27). This formulation has been spurred by the development of tools like multi-slice community detection (28), which can be used to infer activated circuits and their reconfiguration from neuroimaging data collected as participants perform cognitively demanding tasks (29–31). These tools have recently been leveraged to understand the role of dynamic brain-wide connectivity in motor skill learning (29, 30, 32). A key measure from this field is an index of a brain region's tendency to communicate with different networks over time, known as “flexibility” (29, 31). Prior work has shown that flexibility across a number of brain regions predicts individual differences in the speed of acquisition of a simple motor task (29). Network flexibility has also been shown to correlate with working memory and other dimensions of executive function (31). But its role in updating choice behavior based on reinforcement–an inherently dynamic process–is not known.

Guided by the anatomical and computational considerations outlined above, we hypothesized that temporal network dynamics, indexed by flexibility, support key processes underlying reinforcement learning. Specifically, we reasoned that reinforcement learning is associated with dynamic coupling between areas of the striatum and cortical regions processing sensory and value information. We hypothesized that (1) reinforcement learning involves flexible network coupling between the striatum and distributed brain circuits; and (2) that these circuit changes relate to measurable changes in behavior, specifically *learning performance* (accuracy, within subjects) as well as *learning rate* and *inverse temperature*, individual difference measures derived from reinforcement learning models. We were particularly interested in learning rate because it quantifies the extent to which a learner weighs reinforcement from individual trials to update responses. Thus, learning rate is a good index of integration across trials: lower learning rates indicate that value is updated over the course of many trials. Inverse temperature measures how strongly an individual utilizes this learned value in decision-making.

Our final hypothesis was concerned with the relationship between network flexibility and a distinct form of learning: episodic memory for individual events. The rationale for testing episodic memory was two-fold. First, it provided a control comparison for time-on-task effects. Second, it was a question of interest given that very little is known about how episodic memory is supported by network dynamics. Given the extensive literature indicating that separate brain regions support episodic memory vs. reinforcement learning (33, 34), we hypothesized (3) that a distinct set of regions would exhibit a relationship between network flexibility and episodic memory.

To test these hypotheses, we used functional magnetic resonance imaging (fMRI) to measure changes in brain network structure while participants engaged in a reinforcement learning task (**Fig. 1A**). On each trial, participants were presented with a visual cue, made a choice indicated by a key press, and then received feedback. We used a task for which behavior has been well described by reinforcement learning models (35), and which is known from fMRI to involve the striatum (35) and from patient studies to depend on it (34). The task also included trial-unique images presented during feedback, allowing us to test the role of network dynamics in episodic memory. Each of these images coincided with reinforcement, but they were incidental to the learning task (**Fig. 1B**).

**Figure 1.**
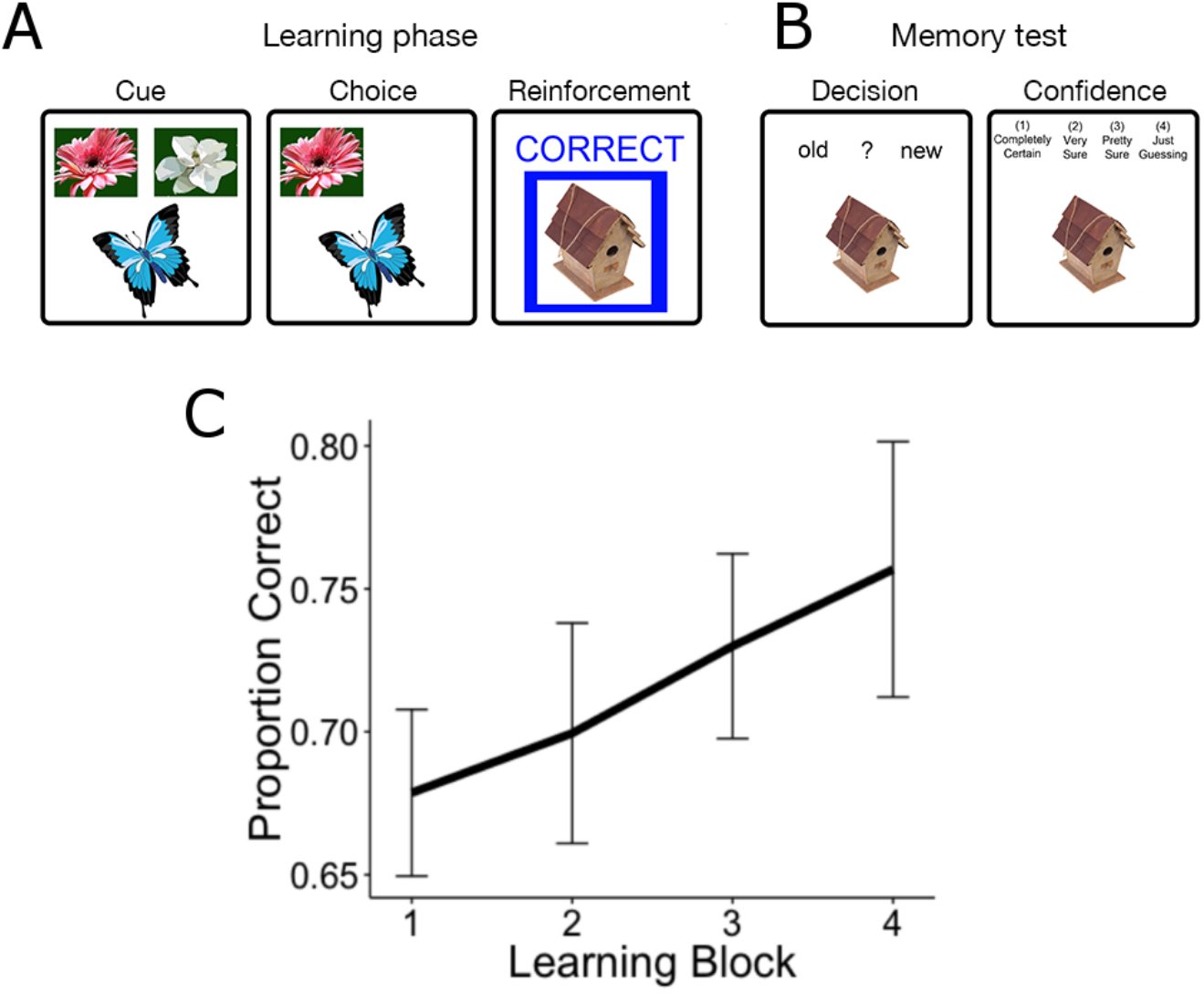
Task design and learning performance. Participants performed a reinforcement learning task while undergoing fMRI (35). **A. Learning phase.** Participants were instructed to associate each of 4 cues (butterflies) with one of two outcomes (flowers). Feedback was probabilistic, with positive feedback following the choice on 80% of correct trials and on 20% of incorrect trials. **B. Memory test.** Each feedback event was presented with a unique image. Thirty minutes following the MRI scan, participants were given a surprise episodic memory test, testing recognition and confidence for images seen during the scan, intermixed with novel images. **C.** Average performance on the learning task improved linearly, suggesting continuous learning across all trials.

## Results

### Reinforcement learning performance

Participants learned the correct response for each cue. The percentage of optimal responses increased continuously from 68% in the first block to 76% in the final block, on average. Using a mixed-effects logistic model, we observed a significant effect of block on learning performance, as measured by the proportion of optimal responses during each block (**Fig. 1C**; β = 0.28, Standard Error (S.E.) = 0.11, p = 0.01 (Wald approximation, (36)). We also fit reinforcement learning models (1, 5) to participants’ trial-by-trial choice behavior, utilizing hierarchical Bayesian models to aid estimation and pool information across subjects ((37), **SI**).

Of particular interest was the learning rate *α*, a parameter that indexes the extent to which an individual weighs feedback from single trials. A low learning rate indicates that an individual is combining choice value over multiple experiences. Also of interest was inverse temperature β, which measures how strongly an individual relies on learned value overall. The average learning rate (*α*) was 0.41 with a standard deviation of 0.14; the average inverse temperature (β) was 3.84, with a standard deviation of 4.31 (see **SI** for details about models and fit). These *α* and β parameters provide a mechanistic probe of individual differences in learning, which allowed us to characterize the relationship between network dynamics and distinct sources of learning variability.

### Flexibility in the striatum relates to reinforcement learning

As a first step, we sought to characterize spatial and temporal properties of dynamic brain networks during the task. We constructed dynamic functional connectivity networks for each subject in 50 s windows, and used a recently developed multi-slice community detection algorithm (28) to partition each network into dynamic communities: groups of densely connected brain regions that evolve in time (**SI**). Our analyses included 110 cortical and subcortical ROIs from the Harvard-Oxford atlas, including bilateral nucleus accumbens, caudate, and putamen subregions of the striatum. We computed a flexibility statistic for each learning block, which measures the proportion of changes in each region's allegiance to large-scale communities over time (29).

To test whether flexibility in the striatum's network coupling is related to learning performance, we fit a mixed effects logistic regression (36) using average flexibility across the Harvard Oxford striatum ROIs during individual learning blocks to predict performance. Striatal flexibility computed for each block was significantly associated with the proportion of optimal responses (**Fig. 2A**; β = 9.45, S.E. = 2.75, p < 0.001 (Wald approximation (36))). This effect could not be distinguished statistically across subregions of the striatum (**Fig. 2B**, **SI**). For appropriate posterior inference, we fit a Bayesian extension of this model (38) (**SI**), to generate a posterior 95% credible interval of [3.53, 14.99] (**Fig. S4**). To ensure that this approach reflected a within-subjects relationship between flexibility and learning, we included each subject's average flexibility across blocks in the model. This model produced similar results (β = 9.*79*, S.E. = 2.82, p = 0.0005), indicating that increases in dynamic striatal connectivity are associated with increased reinforcement-learning performance.

**Figure 2.**
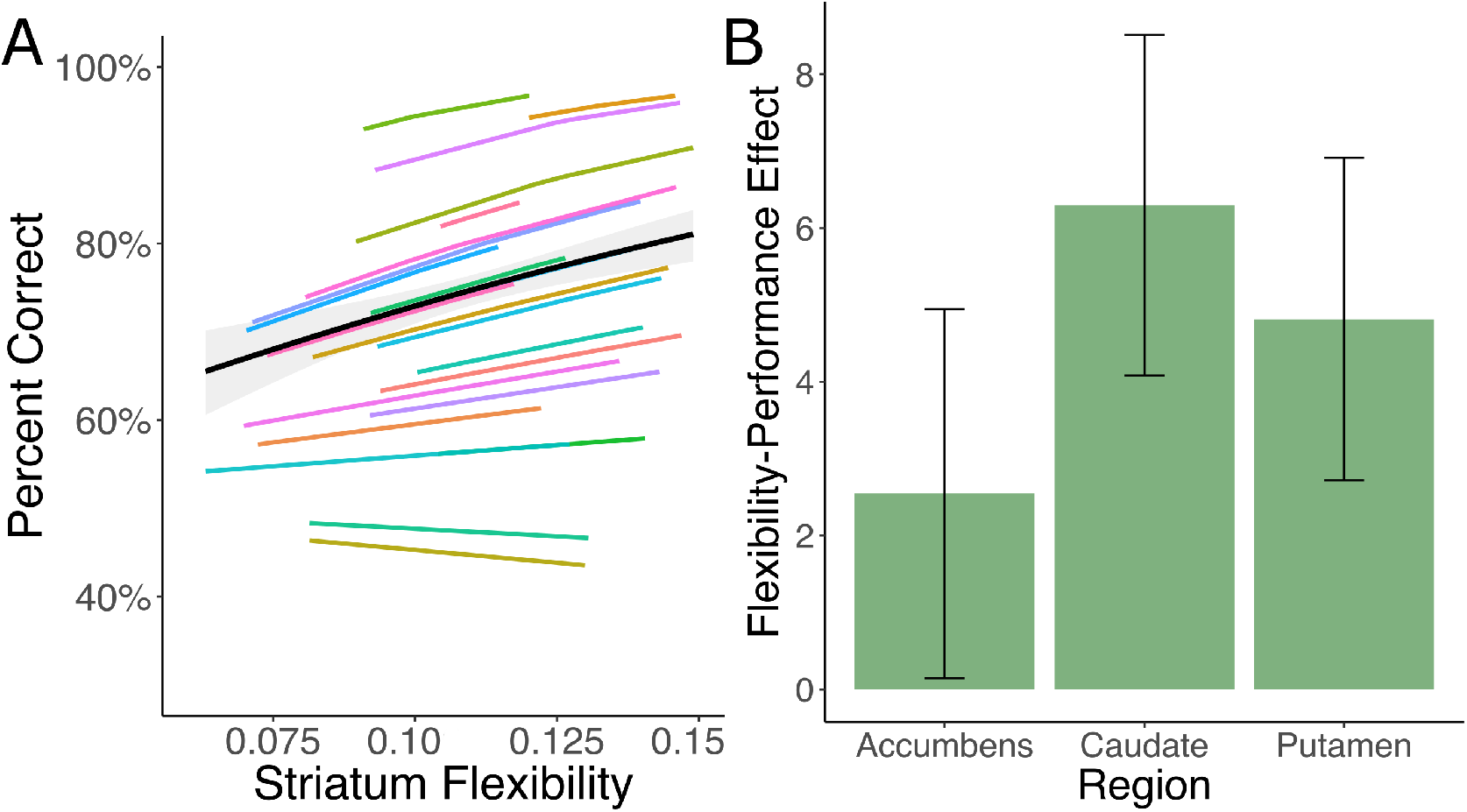
Flexibility in the striatum relates to learning performance within subjects. **A.** Mixed-effects model fit for the association between network flexibility in an *a priori* striatum ROI and learning performance. The black line represents the fixed effect estimate and the gray band represents the 95% confidence interval for this estimate, while color lines represent subject-level random effects estimates. See **SI** for a Bayesian extension of this model and full model fits with uncertainty for individual subjects. **B.** This effect was not distinguishable across regions of the striatum (**SI**; bar plots and error bars represent means and standard errors).

### Individual differences in reinforcement learning parameters correlate with striatal flexibility

We next explored the relationship between flexibility and reinforcement learning model parameters, which account for individual differences in learning behavior. We were most interested in the learning rate *α*, which quantifies the extent to which individuals weigh feedback from single trials when updating the value of a choice (1, 5). Learning rate was negatively correlated with network flexibility in the nucleus accumbens (Spearman's correlation coefficient *ρ* = −0.29, p(*ρ*>0) = 0.04, **Fig. 3A**) and to a lesser extent the caudate (*ρ* = −0.24, p(*ρ*>0) = 0.09 **Fig. 3A**); that is, participants with a lower learning rate (indicating more integration of value across multiple trials) had more flexibility in these regions. Inverse temperature was positively correlated with flexibility in the same regions (accumbens *ρ* = 0.30, p(*ρ*<0) < 0.001; caudate *ρ* = 0.33, p(*ρ*<0) = 0.002), indicating that subjects relying more on learned value overall showed more dynamic striatal connectivity (**Fig. 3B**). The effects of these two parameters were separable, as indicated by partial correlations and the joint posterior distribution (**Fig. S5**).

**Figure 3.**
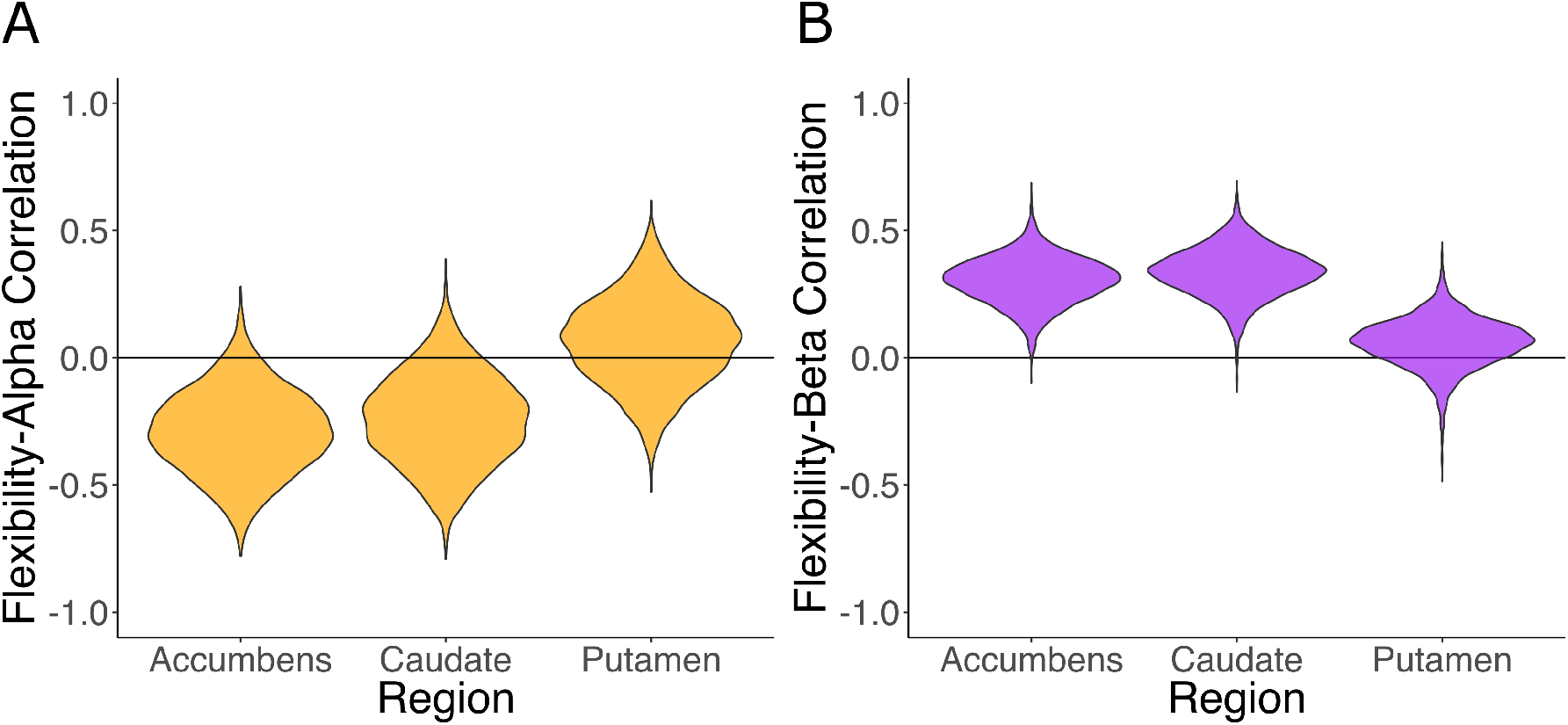
Flexibility in the striatum relates to reinforcement learning parameters across subjects. Violin plots showing posterior distributions of the correlation between parameters from RL models and flexibility in striatal regions. **A**. Learning rate, which indexes reliance on single trials for updating value, is negatively correlated with flexibility in the nucleus accumbens and caudate. **B.** Inverse temperature, which measures overall use of learned value, is positively correlated with flexibility in the same regions. Plotting the joint distribution and utilizing partial correlations indicate that these effects are separable (**Fig. S5**).

Together, these results demonstrate that reinforcement learning involves dynamic changes in network structure centered on the striatum. They also suggest that distinct sources of individual differences in learning–reliance on individual trial feedback and overall use of learned value–are related to differences in this dynamic striatal coupling. We next sought to examine *which* regions the striatum connects with during the task, and how such connections change over the course of learning.

### Striatal allegiance with visual and value regions increases during learning

While an increase in flexible striatal network coupling is associated with learning within and between individuals, this leaves open the critical question of which regions are involved in this process. To address this question we used a dynamic graph theory metric known as *module allegiance*, which measures the extent to which each pair of regions shares a common network during a given time window (30). Using the community labels described above, we first estimated the allegiance between the striatal subregions and every other ROI in the brain for each time window, and then probed their relationship to learning.

We found that overall, the nucleus accumbens and the caudate showed stronger connectivity with midline prefrontal, temporal, and retrosplenial structures, while the putamen exhibited relatively stronger connectivity with motor cortices (**Fig. S6**). To address the key question of which regions changed coupling with subregions of the striatum during learning, we ran whole-brain searches of separate mixed-effects ANOVAs for each region, predicting striatal allegiance with learning block. This analysis does not assume any shape or direction to these temporal changes. A number of regions of visual cortex showed an increase in striatal allegiance over the course of learning (all FDR p<0.001, **Fig. 4**). Examining subregions of the striatum (correcting for multiple comparisons across allegiance of all ROIs with all striatal regions) revealed increases in visual coupling in the nucleus accumbens and putamen, as well as between the putamen and orbitofrontal and ventromedial prefrontal cortices, regions known for their role in value processing (39)(**Fig. S7**).

**Figure 4.**
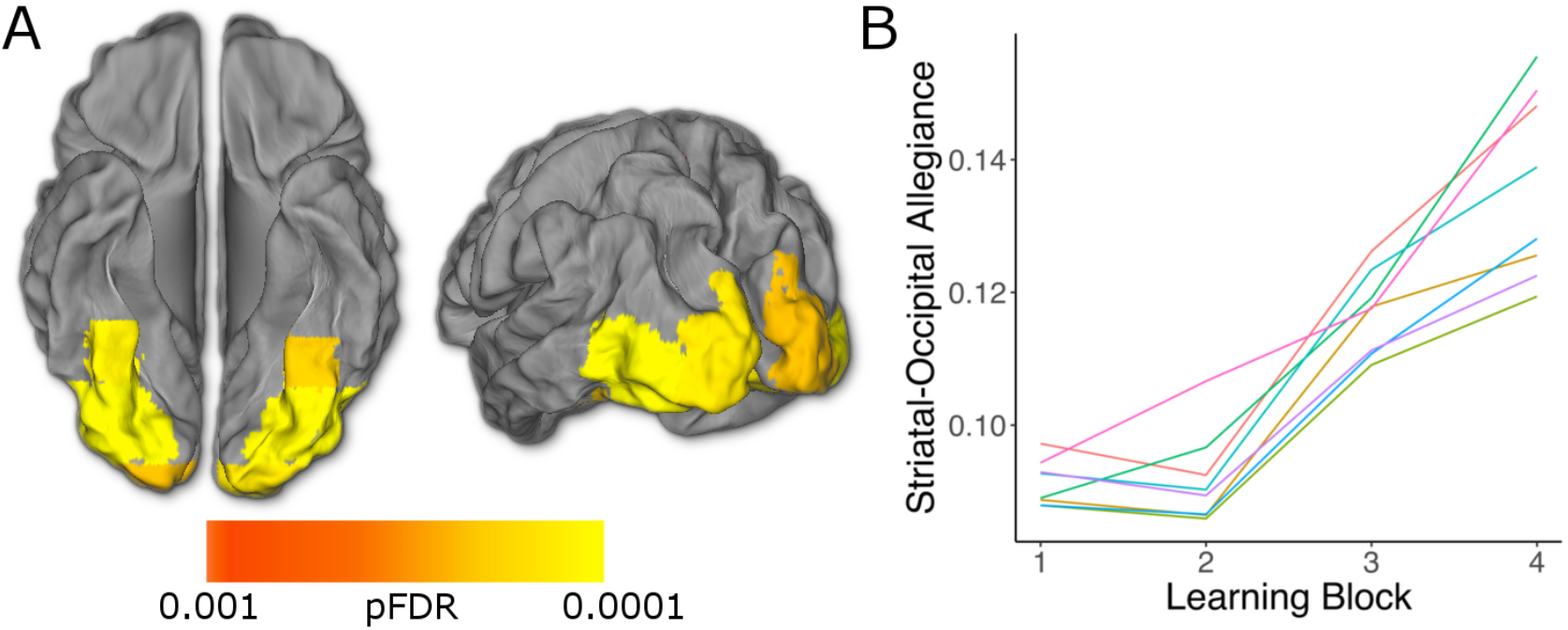
Allegiance between the striatum and visual cortex increases over the course of learning. **A.** Module allegiance between the striatum and a number of visual cortex ROIs changes over time (whole-brain corrected, pFDR < 0.05). **B.** Striatal allegiance increases in each of these visual ROIs (color lines represent the mean for each ROI passing FDR threshold across subjects and striatal regions). Allegiance is averaged across striatal sub-regions. See **Fig. S7** for results presented by subregion.

### Flexibility relates to learning in a distributed set of brain regions

In a number of reports on dynamic networks, averaged whole brain flexibility has been used as a marker of global processes and associated with cognition (29, 40). Indeed, we found that whole-brain flexibility related to learning performance within subjects (β = 11.84, S.E. = 3.91, p<0.005) and, to some extent, learning rate across subjects (**Fig. S8**, *ρ* = −0.26, p(*ρ*>0) = 0.07). We were thus interested in regions outside of the striatum that exhibit dynamic connectivity related to learning.

We conducted the learning performance analysis for each of the 110 ROIs, in addition the analysis in the striatum reported above (**Fig. 2**). This analysis again revealed a significant effect of flexibility in striatal subregions (the right putamen and left caudate) surviving FDR correction. In addition, the whole-brain corrected results, presented in **Fig. 5**, indicate that network flexibility in regions of the motor cortex, parietal lobe, and orbital frontal cortex (among others, see **Table S1** and **Fig. S9** for full list and uncorrected map), are associated with reinforcement learning.

**Figure 5.**
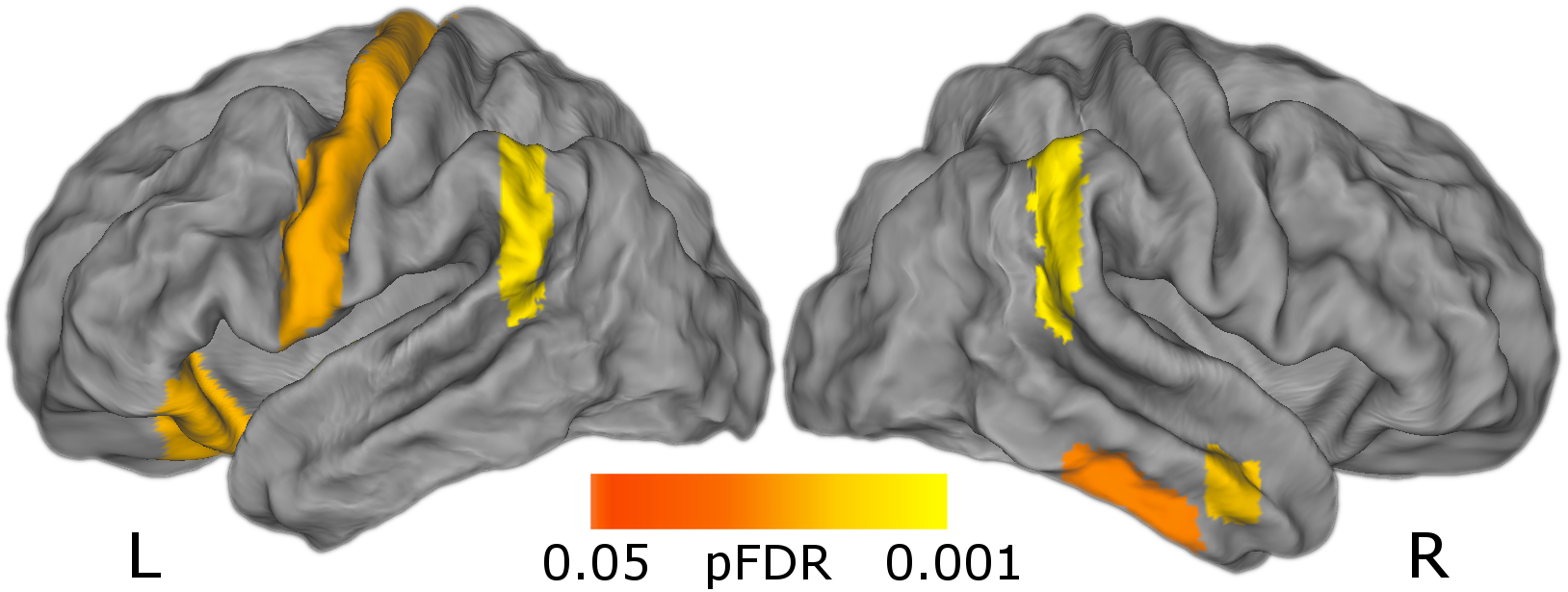
Flexibility in cortical regions is related to learning performance. Regions passing FDR correction following a univariate whole-brain analysis using the same mixed-effects model as the *a priori* striatum ROI. Regions passing this threshold include left motor cortex, bilateral parietal cortex, and right orbitofrontal cortex. See **Table S1** and **Fig. S9** for a full list of regions and an exploratory uncorrected map.

### Flexibility in medial cortical regions is associated with episodic memory

Finally, our task also included trial-unique objects presented simultaneously with reinforcement, allowing us to measure subjects’ episodic memory, a process thought to rely on distinct cognitive and neural mechanisms to feedback-based incremental learning. We tested whether network flexibility was associated with episodic memory for these trial-unique images, as assessed in a later surprise memory test (**Fig. 1**). Having a measure of episodic memory for the same trials in the same participants allowed us to determine whether striatal network dynamics are correlated with any form of learning, or whether these two forms of learning, occurring at the same time, are related to distinct network dynamics.

Behaviorally, participants’ memory was better than chance (d-prime = 0.93, t_21_ = 7.27, p < 0.0001). Memory performance (“hits”) varied across learning blocks, allowing us to assess within-subject associations between network flexibility and behavior (**Fig. 6A**). Memory performance was not correlated with incremental learning performance (mixed effects logistic regression, β = 0.41, Standard Error (S.E.) = 0.51, p = 0.42 (Wald approximation)). We tested the effect of flexibility on memory performance (proportion correct) in each of the 110 ROIs. A whole-brain FDR-corrected analysis revealed one region where flexibility was associated with episodic memory, the left paracingulate gyrus. An exploratory uncorrected analysis revealed regions in the medial prefrontal and medial temporal (parahippocampal) cortices where flexibility was associated with episodic memory (p<0.05 uncorrected, **Fig. 6B**). None of the sub-regions from our *a priori* striatum ROI passed even this low threshold for an effect of flexibility on episodic memory.

**Figure 6.**
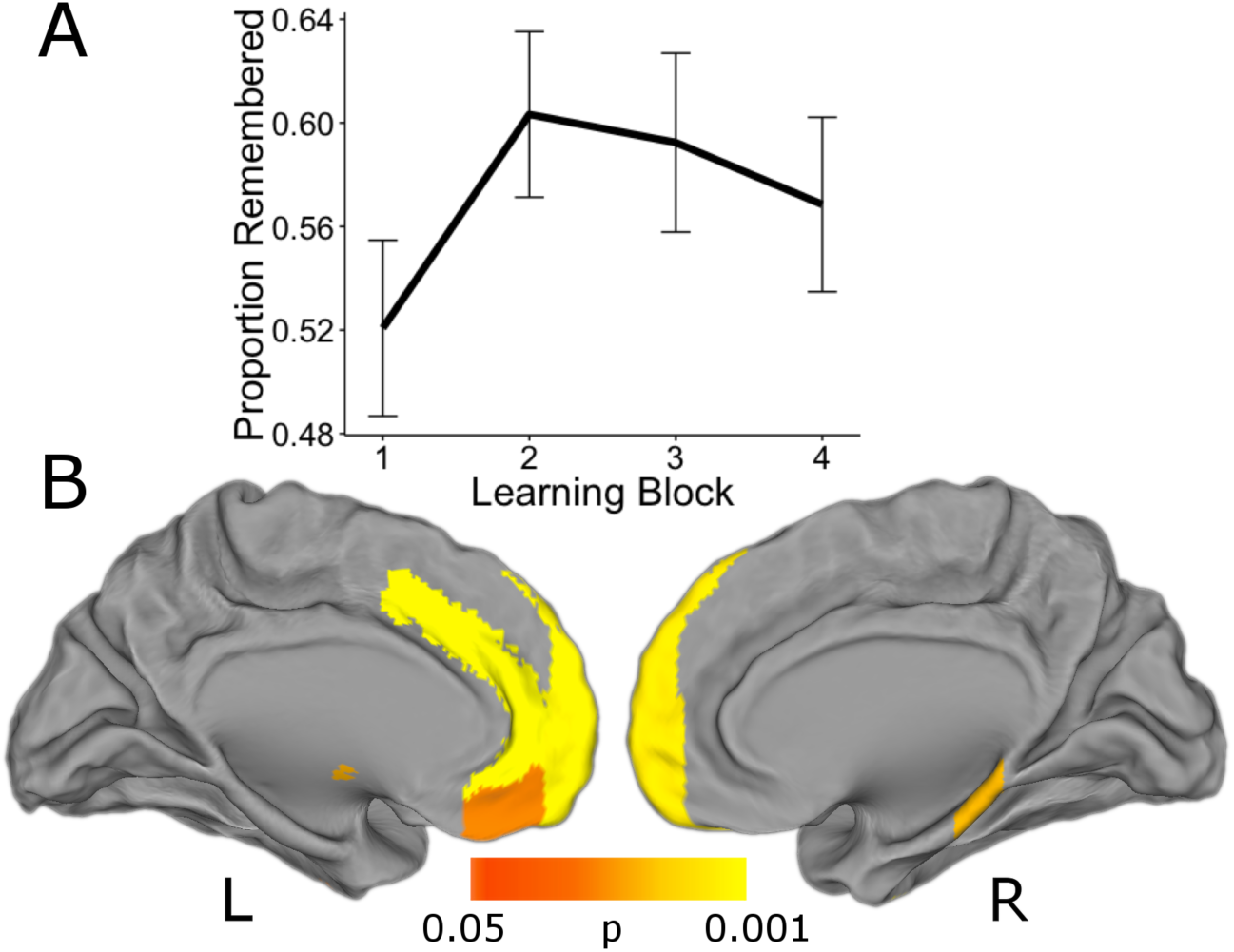
Network flexibility in medial prefrontal and parahippocampal cortex relates to episodic memory. An exploratory analysis showed effects of network flexibility on episodic memory performance in medial prefrontal and temporal lobes. **A**. Average memory (proportion remembered) across blocks. Participants’ recollection accuracy varied across blocks. Line represents group average and bars represent standard errors. **B**. A number of medial prefrontal regions as well as the right parahippocampal gyrus passed an exploratory uncorrected threshold of p<0.05 for the effect of flexibility on subsequent episodic memory. The effect in the left paracingulate gyrus survived FDR correction.

## Discussion

The current study reveals that reinforcement learning involves dynamic coordination of distributed brain regions, particularly interactions between the striatum and visual and value regions in the cortex. Increased dynamic connectivity between the striatum and large-scale circuits was associated with learning performance as well as with parameters from reinforcement learning models. Together, these findings suggest that network coordination centered on the striatum underlies the brain's ability to learn to associate values with sensory cues.

Our results indicate that during learning the striatum increases the extent to which it couples with diverse brain networks, specifically with regions processing value and relevant sensory information. This may represent the formation of efficient circuits for integrating and routing decision variables. Our reinforcement learning model findings are consistent with this idea. Striatal flexibility is negatively related to learning rate, suggesting that increased dynamic coupling with relevant cortical areas may lead to less trial-level weighting of prediction errors during learning. Flexibility in striatal circuits is also positively related to inverse temperature, indicating that this increased dynamic coupling is associated with stronger reliance on learned value during decision-making. Thus, our findings support a framework wherein network flexibility underlies information integration during learning.

This framework offers clear and testable predictions for future studies. It suggests that flexibility will play a larger role in learning the more that learning depends on widespread information integration, and also that this process is specific to regions known to support the particular demands of learning in a given situation. For example, instrumental conditioning involving complex audio-visual stimuli (41) would be expected to associate more strongly with striatal flexibility than the task presented here and would be expected to involve increases in striatal interactions with auditory as well as visual cortex. In addition, learning that relies on other forms of integration, for example the transfer or generalization of information (42, 43) or encoding of associations across space or time (44, 45), is predicted to be associated with network flexibility in medial temporal and prefrontal regions.

There are a number of limitations to this report. First, given the static feedback probabilities, the relationship between network flexibility and learning performance could be affected by the time on task for each subject. This seems unlikely to fully explain the relationship because flexibility was related to multiple aspects of learning behavior and episodic memory, which was associated with network flexibility in a distinct set of brain regions, did not increase over time. Nonetheless, future studies incorporating reversal periods to dissociate time from performance will be important for addressing this issue. Another limitation is the hard-partitioning approach for network assignments provided by multi-slice community detection, which necessarily underemphasizes uncertainty about community labels. There have been very recent attempts to formalize probabilistic models of dynamic community structure (46, 47), but most work examining dynamic networks in the brain have used deterministic community assignment (29, 30, 48). More work is needed to develop and validate these probabilistic models and apply them to neuroscience data. Finally, while the spatial resolution of fMRI makes it an appealing method to characterize dynamic networks, studies using modalities with higher temporal resolution such as ECoG (49) and MEG (50) will be important for providing more fine-grained temporal information.

To summarize, we report a novel link between reinforcement learning and dynamic changes in networks centered on the striatum. While most descriptions of reinforcement learning have focused on the role of individual regions, recent advances in network theory are beginning to make the role of dynamic communication between individual regions and broader networks in this process a tractable area of research (27, 29–31). Here we show that incremental learning based on reinforcement is associated with dynamic changes in network structure, across time and across individuals. Our results suggest that the striatum's ability to dynamically alter connectivity with sensory and value-processing regions provides a mechanism for information integration during decision-making and that learning may be characterized by the formation of these dynamic circuits.

## Materials and Methods

### Experimental Design

Twenty-five healthy right-handed adults (age 24-30 years, mean of 27.7, standard deviation of 2.0, 13 females) were recruited from the University of California Los Angeles and the surrounding community as the adult comparison sample in a developmental study of learning (51). All participants provided informed consent in writing to participate in accordance with the UCLA Institutional Review Board, which approved all procedures. Individuals were paid for their participation. Participants reported no history of psychiatric or neurological disorders, or contraindications for MRI scanning. Three subjects were excluded from this analysis (two for technical issues in behavioral data collection and one for an incidental neurological finding), leading to a final sample size of 22.

### Task and Behavioral Analysis

The probabilistic learning task administered to subjects undergoing an fMRI session has been previously described (34, 35, 51, 52). During the imaging session, individuals underwent an instrumental conditioning procedure, in which they learned to associate 4 cues with 2 possible outcomes. The cues were images of butterflies; the choices were images of flowers. They were then given feedback consisting of the words ‘Correct’ or ‘Incorrect’. Presentation of feedback also included an image of an object unique to each trial, shown in random order for the purpose of subsequent memory testing. For each butterfly image, one flower represented the ‘optimal’ choice, with a 0.8 probability of being correct, while the alternative flower had a 0.2 probability of being followed by correct feedback. Subjects performed four blocks of this probabilistic learning phase, each consisting of 30 trials. Feedback was presented for 2 seconds, and was followed by a randomly jittered inter-trial interval.

For each trial in the learning phase, both the feedback received as well as whether or not subjects made the optimal choice were recorded, and percent correct for each block was computed as the percent of trials on which subjects made the optimal choice, regardless of feedback. These variables enable a characterization of learning as the proportion of optimal choices in each block, as well as that in the test phase. Using this information, we fit reinforcement learning models to subjects’ decisions (1, 5), utilizing a hierarchical Bayesian approach to pool uncertainty across subjects and aid in model identifiability (**SI**).

Following the fMRI session (30 minutes), subjects were given a surprise memory test for the trial-unique object images presented during feedback in the learning phase. Subjects were presented with all 120 objects shown during the conditioning phase, along with an equal number of novel objects, and asked to judge the images as “old” or “new”. They were also asked to rate their confidence for each decision on a scale of 1-4 (one being most confident; four indicating “guessing”). All responses rated 4 were excluded from our analyses (35).

### Dynamic Connectivity Analysis

MRI images were acquired on a 3 T Siemens Tim Trio scanner using a 12-channel head coil (EPI TR=2 s, see **SI** for full acquisition parameters). Functional images were preprocessed using FSL's FMRI Expert Analysis Tool (FEAT (53)). To assess dynamic connectivity between the ROIs, time courses were further subdivided into sub-blocks of 25 TRs each. We then computed the pairwise coherence between each pair of ROIs at *f =* 0.06-0.12 Hz to form temporal connectivity matrices (**SI**). Each connectivity matrix is treated as a graph or network, in which each brain region is represented as a network node, and each functional connection between two brain regions is represented as a network edge (20, 54).

### Uncovering Evolving Circuits Using Multi-slice Community Detection

To extract modules or communities from a single-network representation, one typically applies a community detection technique such as modularity maximization (55). However, these single-network algorithms do not allow for the linking of communities across time-points, thus hampering statistically robust inference regarding the reconfiguration of communities as the system evolves (28). In contrast, the multilayer approaches allow for the characterization of multi-layer network modularity, with layers representing time windows. In this framework, each network node in the multi-layer network is connected to itself in the preceding and following time windows in order to link networks in time. This enables us to solve the community-matching problem explicitly within the model (28), and also facilitates the examination of module reconfiguration across multiple temporal resolutions of system dynamics (56). We thus constructed multilayer networks for each subject, allowing for the partitioning of each network into communities or modules whose identity is robustly tracked across time windows (**SI**). We used the community labels to compute flexibility and module allegiance statistics.

### Dynamic Network Statistics—Flexibility and Allegiance

To characterize the dynamics of these temporal networks and their relation to learning, we computed the *flexibility* of each node, which measures the extent to which a region changed its community allegiance over time (29). Intuitively, flexibility can be thought of as a measure of a region's tendency to communicate with different networks during learning. Flexibility is defined as the number of times a node displays a change in community assignment over time, divided by the number of possible changes (equal here to the number of time windows in a learning block minus 1). This was computed for each region in each block (**Fig. S3**). In addition, average measures of flexibility were computed across the brain and across all blocks. We also computed the *module allegiance* of each ROI with respect to regions of the striatum during each learning block. Module allegiance is the proportion of time windows in which a pair of regions is assigned the same community label, and thus tracks which regions are most strongly coupled with each other at a given point in time. To obtain stable estimates, we averaged both flexibility and allegiance scores for each ROI over the 500 iterations of the multilayer community detection algorithm.

### Relating Dynamic Networks to Reinforcement Learning

To examine the effect of flexibility on learning from feedback, we estimated a generalized mixed-effects model predicting optimally correct choices with flexibility estimates for each block with a logistic link function, using the Maximum Likelihood (ML) approximation implemented in the *lme4* package (36). Subjects’ average flexibility in an *a priori* striatum ROI was used to predict the proportion of optimal choices in each learning block. The ROI included bilateral caudate, putamen, and nucleus accumbens regions from the Harvard-Oxford atlas. We included a random effect of subject, allowing for different effects of flexibility on learning for each subject, while constraining these effects with the group average. Average flexibility across sessions was included as a fixed effect in the model in order to ensure that our estimates represented within-subject learning effects. We also estimated this effect for whole-brain flexibility, which has been related to several cognitive functions in previous reports (29, 31).

To provide appropriate posterior inference about the plausible parameter values indicated by our data, and to account for uncertainty about all parameters, we also fit a fully Bayesian extension of the ML approximation described above for the effect of striatal flexibility on learning performance (**Fig. S4**).

To examine the relationship between flexibility and parameters estimated from reinforcement learning models, we tested whether striatal flexibility was correlated with the learning rate *α* and inverse temperature β for each subject, using Spearman's correlation coefficient due to the non-Gaussian distribution of these parameters. To account for joint uncertainty in these parameters at the group and subject level, correlations were computed over the full posterior distributions from the reinforcement-learning model (**Fig. S1**, **S5**).

To determine which regions changed coupling with the striatum during the course of the task, we fit mixed effects models using learning block to predict log-transformed module allegiance. This analysis was computed first using the average of each ROI's allegiance with the striatal sub-regions, and then separately for all sub-regions of the striatum. In both cases this analysis was carried out for each region of the brain, treating block as a factor so as to avoid assumptions about the linearity or direction of changes in allegiance. We controlled the false discovery rate across all ROI-striatum pairs.

To explore other regions exhibiting effects of dynamic connectivity on learning performance, we separately modeled the effect of flexibility in each brain region on reinforcement learning using the ML approximation implemented in the *lme4* package. We applied a false discovery rate correction for multiple comparisons across regions (57). While regions passing this threshold are reported, we also visualize the results using an exploratory uncorrected threshold of p<0.05 (**Fig. S9**).

To explore the relationship between network dynamics and other forms of learning, we also regressed flexibility statistics from each ROI against subsequent memory scores for the trial-unique objects presented during feedback. If the effects of striatal flexibility were relatively selective to incremental learning, we expected to find no significant association even at an uncorrected threshold with memory in the regions comprising our striatal ROI. In addition, this provided an exploratory analysis to examine the regions in which network flexibility plays a potential role in episodic memory. Given a host of previous studies on multiple learning systems, we reasoned it might be possible to detect an effect of dynamic network coupling on episodic memory in regions traditionally associated with this form of learning.

## Supporting Information

### Task and Behavioral Analysis

Before scanning, participants completed a practice round of 8 trials to become familiar with the task. On each trial, participants were presented with an image of one of the four butterflies along with two flowers, and asked to indicate which flower the butterfly was likely to feed from, using a left or right button press. The four learning blocks were followed by a test phase, in which subjects performed the same butterfly task without feedback for 32 trials.

### Reinforcement Learning Model

To characterize learning, we fit standard reinforcement learning models to individuals’ choice behavior (1, 5). Briefly, the expected value for a given choice at time *t*, Qt, is updated based on the reinforcement outcome r*_t_* via a prediction error *δ*_*t*_:

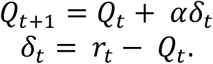

The reinforcement learning models included two free parameters, *α* and β. The learning rate *α* is a parameter between 0 and 1 that measures the extent to which value is updated by feedback from a single trial. Higher *α* indicates more rapid updating based on few trials and lower *α* indicates slower updating based on more trials. Another parameter fit to each subject is the inverse temperature parameter β, which determines the probability of making a particular choice using a softmax function (1, 58), so that the probability of choosing choice 1 on trial *t* would be:

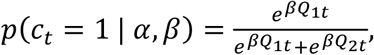

where *p* (*c_t_* = 1) refers to the probability of choice one and *Q*_1*t*_ is the value for this choice on trial *t*.

Reinforcement learning models of this form have known issues with identifiability (37). To constrain the parameter space to reduce noise, we fit a hierarchical Bayesian model, which regularizes this estimation with empirical prior distributions on *α* and β (1):

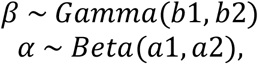

where b1 and a1 are shape parameters, and b2 and a2 are scale parameters. By fitting prior parameters as part of the model, individual-level likelihood parameters are constrained by group average distributions. These group parameters were themselves regularized by weakly informative hyperprior distributions (*Cauchy*^+^ (0, 5) in all cases). Models were fit using Hamiltonian Markov Chain Monte Carlo in Stan (38). In addition to the benefit of constraining the parameter space, this approach produces a posterior distribution of all parameters, which incorporates uncertainty at both group and individual levels in parameter estimation, and also allows for the consideration of all plausible values of RL parameters in subsequent analyses (see **Fig. S1** for examples of group- and subject-level distributions), rather than relying on point estimates or Gaussian assumptions.

### MRI Acquisition Parameters

For each block of the learning phase of the conditioning task, we acquired 200 interleaved T2*-weighted echo-planar (EPI) volumes with the following sequence parameters: TR = 2000 ms; TE = 30 ms; flip angle (FA) = 90°; array =64 × 64; 34 slices; effective voxel resolution = 3×3×4 mm; FOV = 192 mm). A high resolution T1-weighted MPRAGE image was acquired for registration purposes (TR = 2170 ms, TE = 4.33 ms, FA = 7°, array = 256 × 256, 160 slices, voxel resolution = 1 mm^3^, FOV = 256).

### fMRI Preprocessing

Images from each learning block were high-pass filtered at *f* > 0.008 Hz, spatially smoothed with a 5mm FWHM Gaussian kernel, grand-mean scaled, and motion corrected to their median image using an affine transformation with tri-linear interpolation. The first three images were removed to account for saturation effects. Functional and anatomical images were skull-stripped using FSL's Brain Extraction Tool. Functional images from each block were co-registered to subject's anatomical images and non-linearly transformed to a standard template (T1 Montreal Neurological Institute template, voxel dimensions 2 mm^3^) using FNIRT (59). Following image registration, time courses were extracted for each block from 110 cortical and subcortical regions of interest (ROIs) segmented from FSL's Harvard-Oxford Atlas. Due to known effects of motion on measures of functional connectivity (60, 61), time courses were further preprocessed via a nuisance regression. This regression included the six translation and rotation parameters from the motion correction transformation, average CSF, white matter, and whole brain time courses, as well as the first derivatives, squares, and squared derivatives of each of these confound predictors (62).

### Dynamic Connectivity Analysis

For each 25 TR sub-block, connectivity was quantified as the magnitude-squared coherence between each pair of ROIs at *f =* 0.06-0.12 Hz in order to later assess modularity over short time windows in a manner consistent with previous reports (29, 32):

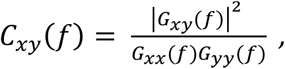

where *G_xy_* (*f*) is the cross-spectral density between regions x and y, and *G_xx_* (*f*) and *G_yy_* (*f*) are the autospectral densities of signals x and y, respectively. We thus created subject-specific 110 × 110 × 32 connectivity matrices for 110 regions and 8 time windows for each of the 4 learning blocks, containing coherence values ranging between 0 and 1. The frequency range of 0.06-0.12 Hz was chosen to approximate the frequency envelope of the hemodynamic response, allowing us to detect changes as slow as 3 cycles per window with a 2 second TR.

In the context of dynamic functional connectivity matrices, the network representation is a *temporal network*, which is an ensemble of graphs that are ordered in time (63). If the temporal network contains the same nodes in each graph, then the network is said to be a multilayer network where each layer represents a different time window (64). The study of topological structure in multilayer networks has been the topic of considerable study in recent years, and many graph metrics and statistics have been extended from the single-network representation to the multilayer network representation. Perhaps one of the single most powerful features of these extensions has been the definition of so-called *identity links*, a new type of edge that links one node in one time slice to itself in the next time slice. These identity links hard code node identity throughout time, and facilitate mathematical extensions and statistical inference in cases that had previously remained challenging.

### Multi-slice Community Detection

While many statistics are available to the researcher to characterize network organization in temporal and multilayer networks, it is not entirely clear that all of these statistics are equally valuable in inferring neurophysiologically relevant processes and phenomena (23). Indeed, many of these statistics are difficult to interpret in the context of neuroimaging data, leading to confusion in the wider literature. A striking contrast to these difficulties lies in the graph-based notion of modularity or community structure (55), which describes the clustering of nodes into densely interconnected groups that are referred to as *modules* or *communities* (65, 66). Recent and convergent evidence demonstrates that these modules can be extracted from rest and task-based fMRI data (67, 68), demonstrate strong correspondence to known cognitive systems (including default mode, fronto-parietal, cingulo-opercular, salience, visual, auditory, motor, dorsal attention, ventral attention, and subcortical systems (22, 69)), and display non-trivial re-arrangements during motor skill acquisition (29, 30) and memory processing (31). These studies support the utility of module-based analyses in the examination of higher order cognitive processes in functional neuroimaging data.

The partitioning of these multilayer networks into temporally linked communities was carried out using a Louvain-like locally greedy algorithm for multilayer modularity optimization (28, 70).

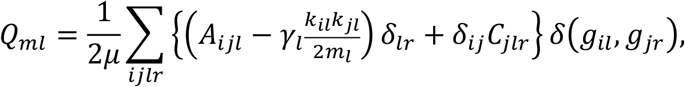

where *Q_ml_* is the multilayer modularity index. The adjacency matrix for each layer *l* consists of components A*_ijl_*. The variable γ*_l_* represents the resolution parameter for layer *l*, while C*jlr* gives the coupling strength between node *j* at layers *l* and *r* (see below for details of fitting these two parameters). The variables *g_il_* and *g_jr_* correspond to the community labels for node *i* at layer *l* and node *j* at layer *r*, respectively; k*_il_* is the connection strength (in this case, coherence) of node *i* in layer *l*; 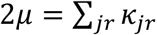; the multilayer node strength *k_jl_* = *k_jl_* + *c_jl_*; and 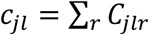. Finally, the function (*g_il_*, *g_jr_*) refers to the Kronecker delta function, which equals 1 if *g_il_*=*g_jr_*, and 0 otherwise.

Resolution and coupling parameters (γ*_l_* and *C_jlr_*, respectively) were selected using a grid search formulated explicitly to optimize *Q_ml_* relative to a temporal null model (56). The temporal null model we employed is one in which the order of time windows in the multilayer network was permuted uniformly at random. Thus, we performed a grid search to identify the values of γ_l_ and *C_jlr_* that maximized *Q_ml_*–*Q_null_* (**Fig. S2**), following (56). To ensure statistical robustness, we repeated this grid search 10 times. To maximize the stability of resolution and coupling, each subject's parameters were treated as random effects, with the best estimate of resolution and coupling generated by averaging across-individual subject estimates. This is a similar approach to that taken in computational modeling of reinforcement learning, in which learning rate and temperature parameters are averaged in order to generate prediction error estimates (1). With this approach, we estimated the optimal resolution parameter *γ* to be 1.18 (standard deviation of 0.61) and the coupling parameter C to be 1 (this was the optimal parameter for all subjects for all iterations). These values are quite similar to those chosen *a priori* (usually setting both parameters to unity) in previous reports (29).

Finally, we note that maximization of the modularity quality function is NP-hard, and the Louvain-like locally greedy algorithm we employ is a computational heuristic with non-deterministic solutions. Due to the well known near-degeneracy of *Q_ml_* (28, 56, 71), we repeated the multi-slice community detection algorithm 500 times using the resolution and coupling parameters estimated from the grid search procedure outlined above. This approach ensured an adequate sampling of the null distribution (56). Each repetition produced a hard partition of nodes into communities as a function of time window: that is, a community or module allegiance identity for each of the 110 brain regions in the multilayer network.

### Flexibility and Learning

Averaged across time, sensory and motor regions showed the lowest levels of flexibility (72), while association cortices showed moderate to high levels of (**Fig. S3a**). In addition to this regional distribution, we sought to verify that flexibility was not related to the size of the ROI. We found that ROI size only explained 1.17% of the variance in average flexibility across individuals, *r* = 0.11, *t*108 = 1.311, *p* = 0.26), indicating that this measure is not an artifact of the parcellation we used. We also studied the temporal profile of this measure, by examining changes in flexibility over the course of the task. Flexibility (averaged across all ROIs) increased in early learning blocks, before slightly but significantly decreasing in later stages of the task (**Fig. S3b**; quadratic effect −0.01; p<0.0001). We fit generalized mixed-effects models of the relationship between learning performance and average striatal flexibility using *lme4*. In an attempt to examine distinct effects in different striatal sub-regions, we attempted to include striatal ROI as a varying effect. The maximum likelihood estimate of the variance by region was 0, indicating that there is little variability across regions and not enough data to distinguish these small effects. **Fig. 2b** shows the results of separate models for each region of the striatum, essentially assuming that this regional variance is infinite. Even with this assumption, there were no significant differences between estimates for any pair of striatal regions.

### Bayesian Models of Flexibility and Performance

We used the ‘brms’ package for fitting flexibility-performance models in the Stan language (38). These were similar to the likelihood approximation models, but included a covariance parameter for subject-level slopes and intercepts (which could not be fit by the above approximation), and weakly informative prior distributions to regularize parameter estimation:

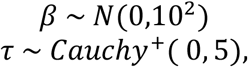

where β represents the “fixed effects” parameters (slope and intercept), *τ* represents the “random effects” variance for subject-level estimates sampled from β, and *Cauchy*^+^ is a positive half-t distribution with one degree of freedom (73). Similarly, we used an lkj prior with *η* = 2 for correlations between subject-level intercept and slope estimates (74). This approach also allowed us to visualize subject-level estimates of the relationship between flexibility and performance (**Fig. S4**).

## Acknowledgements

RTG acknowledges support from the National Institute of Mental Health (F31 MH109247-01A1). DSB acknowledges support from the John D. and Catherine T. MacArthur Foundation, the Alfred P. Sloan Foundation, the Army Research Laboratory and the Army Research Office through contract numbers W911NF-10-2-0022 and W911NF-14-1-0679, the National Institute of Mental Health (2-R01-DC-009209-11), the National Institute of Child Health and Human Development (1R01HD086888-01), the Office of Naval Research, and the National Science Foundation (CRCNS award BCS-1441502 and CAREER award PHY-1554488). AG acknowledges support from the William T. Grant Foundation and the Center for Translational and Prevention Science (P30 DA027827, Brody-PI) funded by NIDA. D.S. acknowledges support from the National Science Foundation (Career Award #0955494), the National Institute of Health (NINDS R01NS079784 and CRCNS R01DA038891), and the McKnight Foundation Memory and Cognitive Disorders Award.

## Supplemental Figures

**Figure S1.**
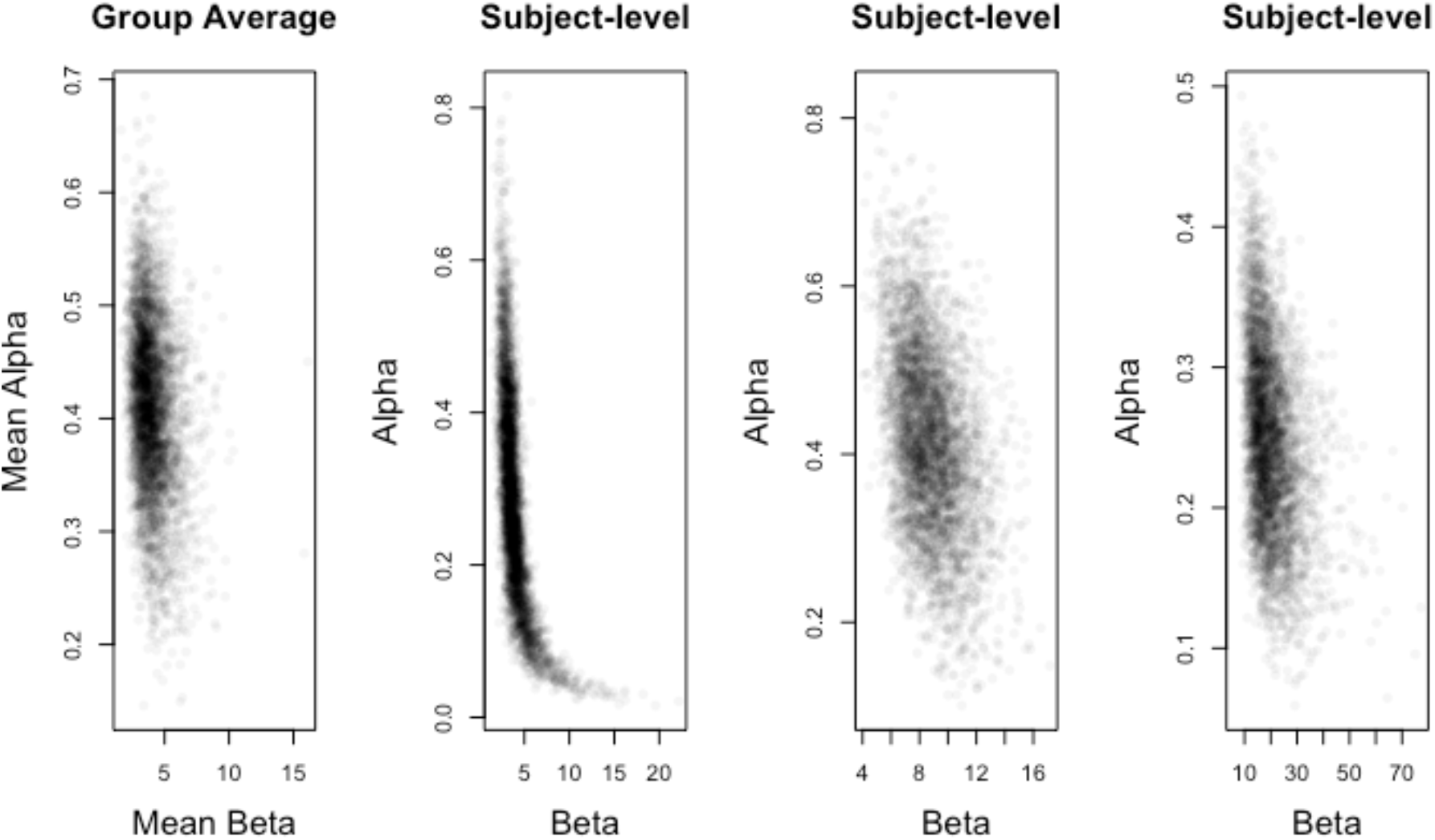
Group- and subject-level estimates of reinforcement learning model parameters. **A.** Joint posterior distribution for group-average learning rates and inverse temperature parameters derived from hierarchical Bayesian reinforcement learning model, each computed from the scale and shape of their respective prior distributions. **B.** Example subject-level parameters from three individuals, exhibiting a range of uncertainty. Note that in individuals with learning rates plausibly close to 0, the inverse temperature becomes highly uncertain for extreme values of learning rate. This uncertainty was accounted for in our correlations with network flexibility by computing these correlations over the full distribution of parameter values.

**Figure S2.**
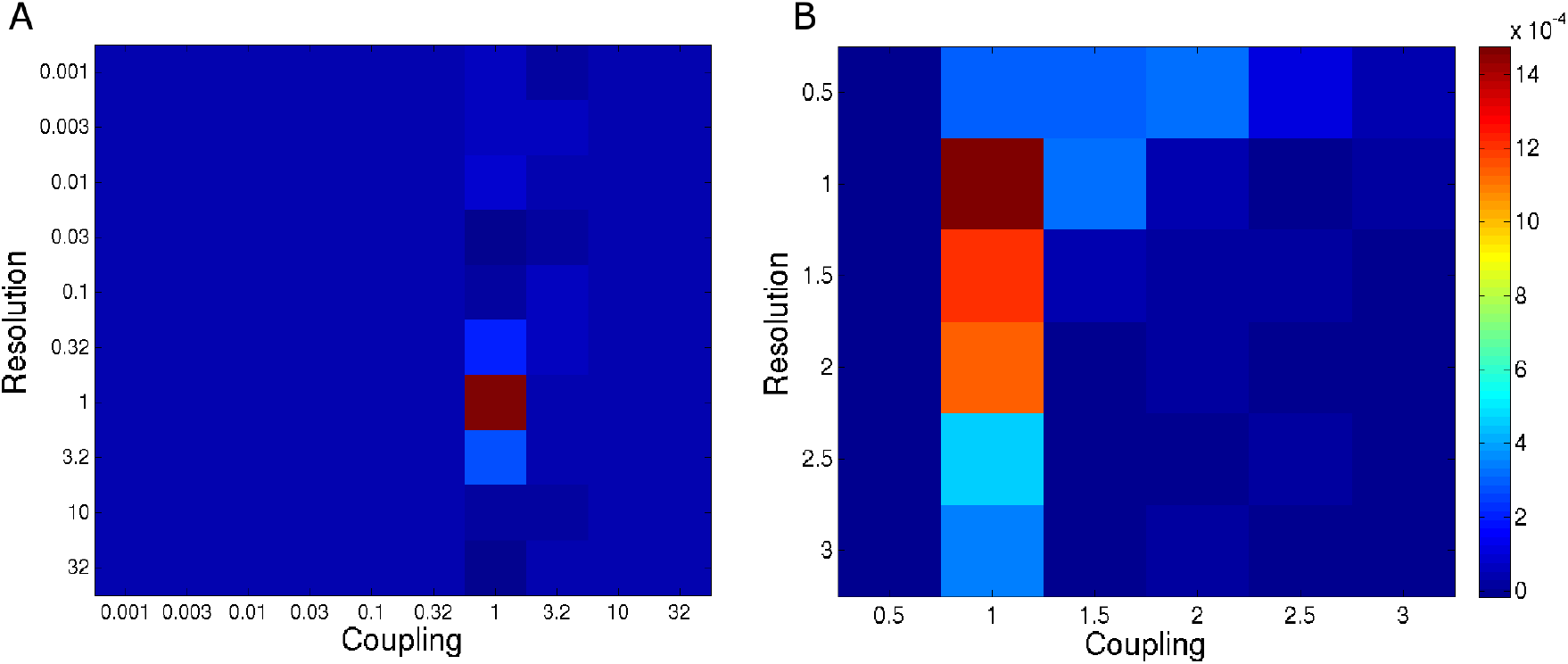
Grid searches for optimizing resolution and coupling parameters for multi-slice community detection. Two grid searches at distinct scales illustrating that our resolution and coupling parameters (1.18 and 1, respectively) fall at the peak of our optimization function. **A.** We first used a wide grid to cover a larger range of potential parameters. The average peak of this search across subjects and iterations was used to select resolution and coupling terms for multi-slice community detection. **B.** To ensure that the large grid steps did not affect this selection, we repeated the search on a smaller scale. Our parameters are clearly within the optimum range on both grids.

**Figure S3.**
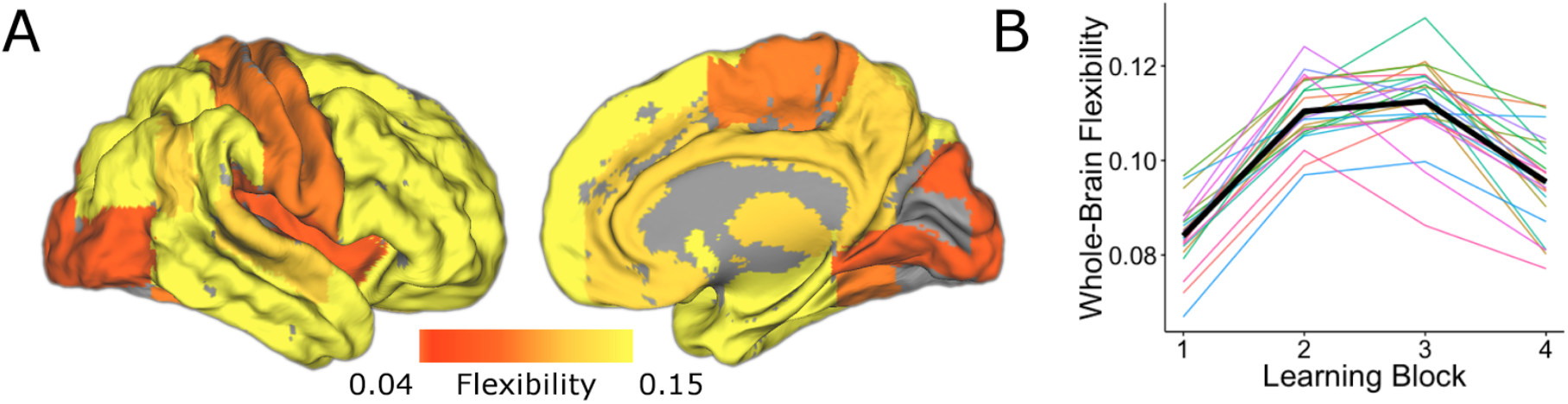
Spatial and temporal characteristics of network flexibility. Network flexibility exhibited distinct spatial and temporal patterns. **A.** Flexibility averaged across all learning blocks was highest across regions of association cortex and lowest in sensory and motor regions. **B.** Flexibility averaged across the whole brain consistently increased in early learning blocks. Color lines represent whole-brain flexibility for single subjects.

**Figure S4.**
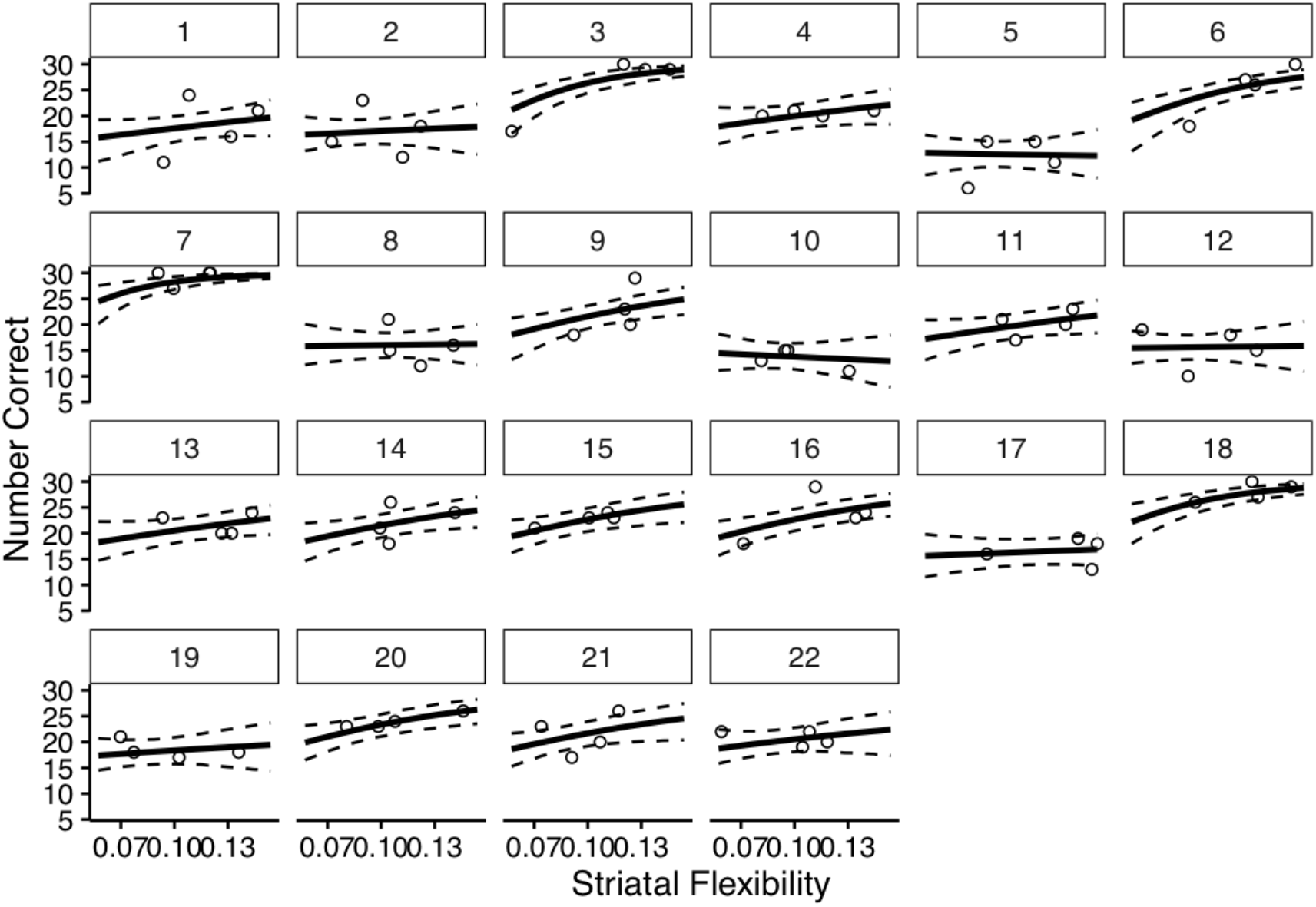
Subject-level data and fits for Bayesian hierarchical model of the effect of striatal flexibility on learning. For posterior inference on the effect of striatal flexibility on learning performance, we fit a Bayesian hierarchical model. Each subplot displays data (open circles) from a single subject. Solid lines represent model estimates for the effect of flexibility on learning, while dotted lines represent 95% credible intervals.

**Figure S5.**
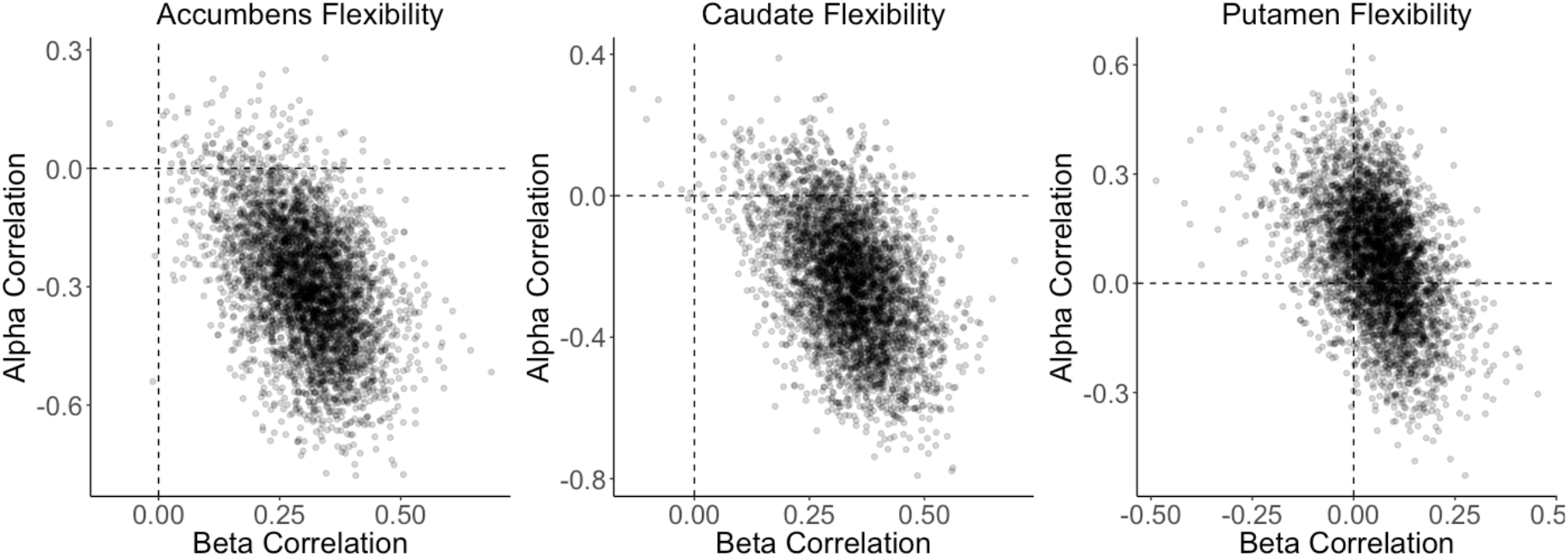
Joint distributions of correlation between flexibility and reinforcement learning model parameters for striatal regions. Because we computed Spearman correlations over the posterior distributions of learning rate and inverse temperature from hierarchical Bayesian models, our inferences can be most fully expressed with the joint distributions of correlations between flexibility and each parameter. While there is some covariance between these correlations, the two effects are clearly separable. This is further supported by partial correlations, which did not substantially alter inference (accumbens-alpha *ρ* = −0.23, caudate-alpha *ρ* = −0.16, accumbens-beta *ρ* = 0.23, caudate-beta *ρ* = 0.28).

**Figure S6.**
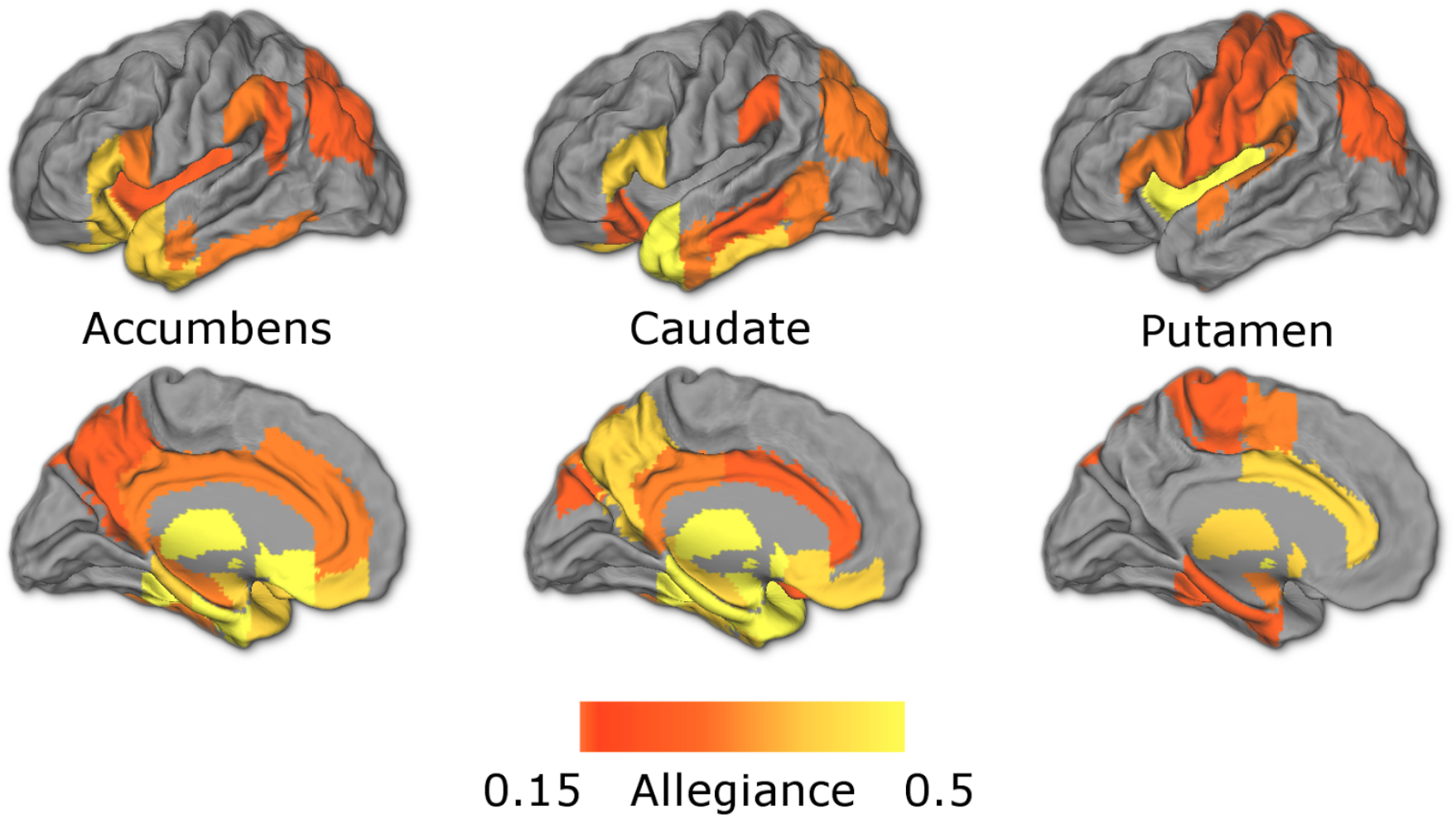
Module allegiance broken down by striatal region. Maps show regions in the 50^th^ percentile of allegiance for each striatal ROI, averaged over all learning blocks. Consistent with anatomical and functional connectivity, the nucleus accumbens and caudate show stronger allegiance with midline frontal, temporal, and retrosplenial regions, while the putamen shows relatively stronger allegiance with motor regions.

**Figure S7.**
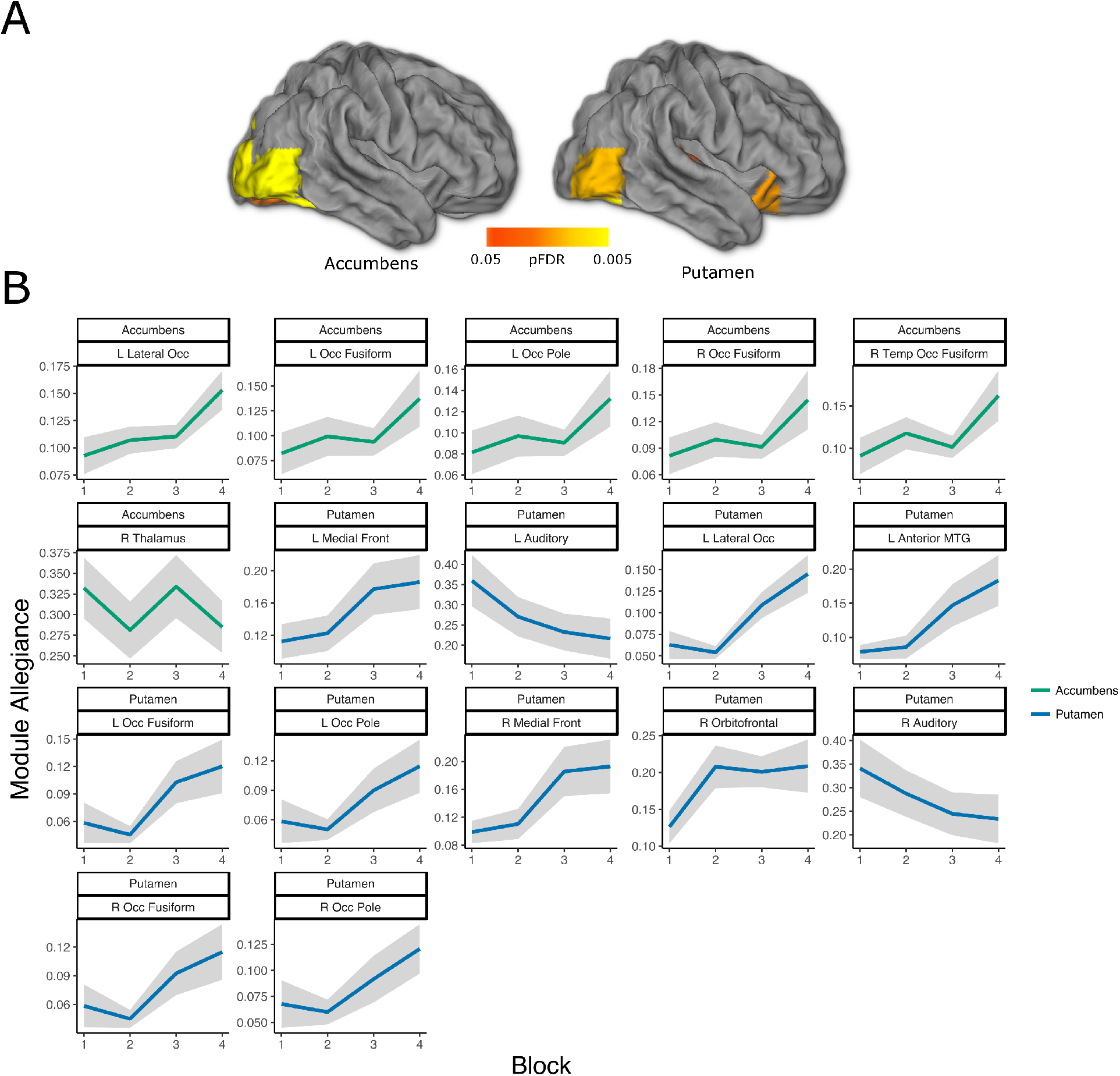
Visual and value regions change allegiance with the striatum over the course of the task. **A.** Regions where time-dependent changes to striatal allegiance exceed a threshold of pFDR<0.05, corrected for all ROIs’ allegiance with all three striatal regions (109 × 3 comparisons). These results were generated using mixed-effects ANOVAs and contain no assumptions about the shape or direction of changes. **B.** Panel plot showing the change in allegiance over time for every pair passing the above threshold. Lines and bands represent and bands standard errors. As with average striatal allegiance, the nucleus accumbens and putamen increase coupling with visual regions during the task. In addition, the putamen exhibits an increase in coupling with the right orbitofrontal and ventromedial prefrontal cortex and a decrease in coupling with primary auditory cortex. No regions’ allegiance with the caudate survived correction for multiple comparisons. Abbreviations: Occ=Occipital, Temp=Temporal, Front=Frontal, and MTG=Middle Temporal Gyrus.

**Figure S8.**
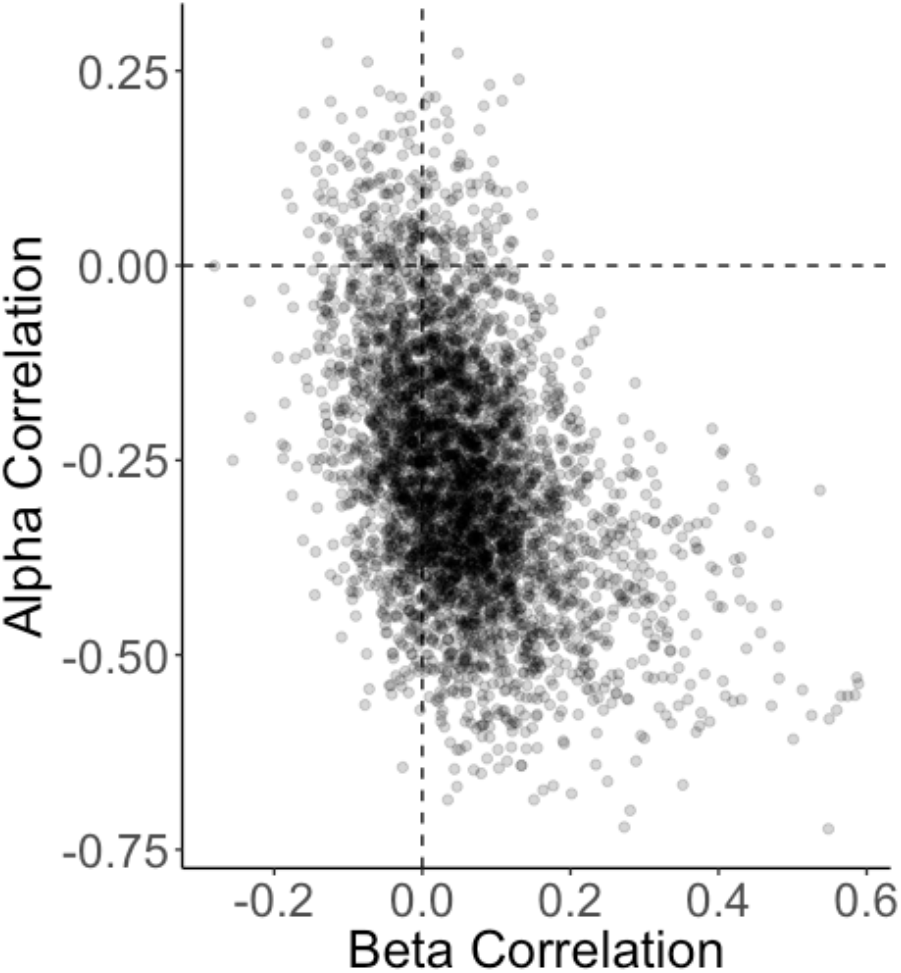
Whole-brain flexibility is negatively correlated with learning rate. Because reinforcement learning parameters were fit using a fully Bayesian model, we computed correlations with flexibility over the full posterior distributions for these parameters. Our uncertainty about these correlations can be expressed with the joint distribution of learning rate and inverse temperature correlations. There was evidence of a negative relationship between whole-brain flexibility and learning rate (*ρ* = −0.26, p(*ρ*>0) = 0.07).

**Figure S9.**
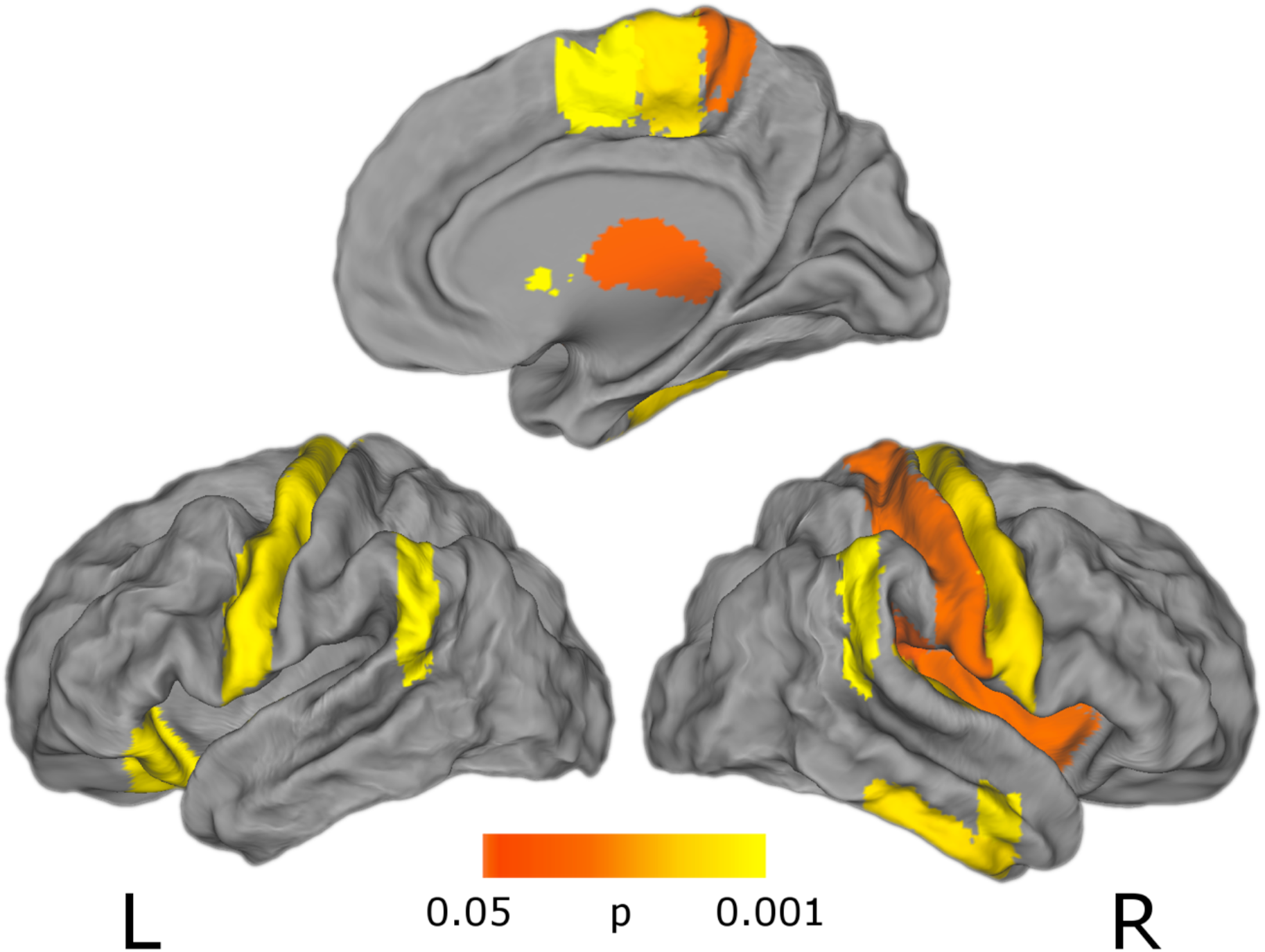
**Exploratory uncorrected (p<0.05) results for whole-brain effect of network flexibility on reinforcement learning performance.**

**Figure S10.**
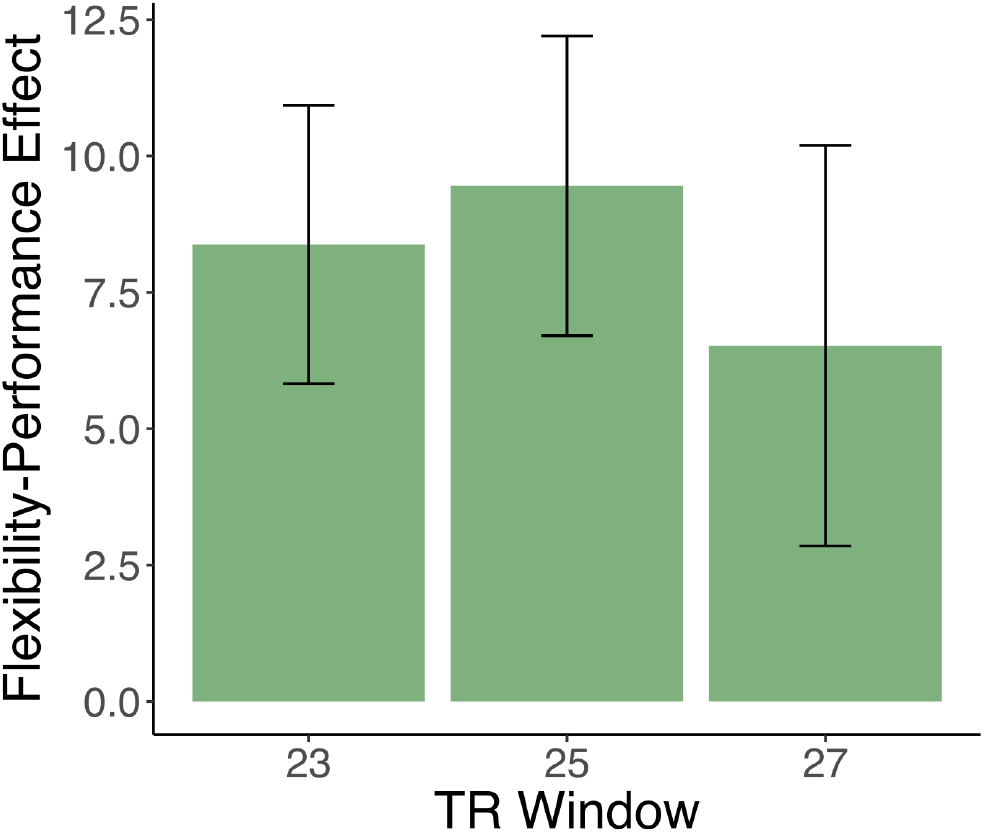
The relationship between striatal flexibility and learning is robust across different lengths of connectivity windows. We recomputed coherence over 23- and 27-TR windows, and ran the multi-slice community detection algorithms to extract flexibility. The relationship between average flexibility and performance was similar across time windows. Regression coefficients and standard errors (SE) from mixed-effects models are plotted here (23 TR: effect=8.38, SE=2.55; 27 TR: effect=6.52, SE=3.67).

**Table S1.**

| Harvard Oxford Region | Regression Coefficient | p |
| --- | --- | --- |
| Left Caudate | 5.93 | 0.003 |
| Left Orbitofrontal Cortex | 4.88 | 0.006 |
| Left Planum Polare | 10.97 | 0.000 |
| Left Precentral Gyrus | 4.79 | 0.006 |
| Left Supramarginal Gyrus | 5.38 | 0.003 |
| Right Inferior Temporal Gyrus, anterior | 4.75 | 0.01 |
| Right Inferior Temporal Gyrus, posterior | 4.66 | 0.01 |
| Right Supplemental Motor Cortex | 8.54 | 0.0009 |
| Right Middle Temporal Gyrus | 3.51 | 0.006 |
| Right Planum Temporal | 8.19 | 0.01 |
| Right Putamen | 6.10 | 0.003 |
| Right Supramarginal Gyrus | 6.48 | 0.003 |

## References

1. Daw ND (2011) Trial-by-trial data analysis using computational models. Decision making, affect, and learning: Attention and performance XXIII 23:3–38.

2. Shohamy D, et al. (2004) Cortico-striatal contributions to feedback-based learning: converging data from neuroimaging and neuropsychology. Brain 127(4):851–859.

3. O'Doherty JP, Dayan P, Friston K, Critchley H, & Dolan RJ (2003) Temporal difference models and reward-related learning in the human brain. Neuron 38(2):329–337.

4. Frank MJ, Seeberger LC, & O'reilly RC (2004) By carrot or by stick: cognitive reinforcement learning in parkinsonism. Science 306(5703):1940–1943.

5. Sutton RS & Barto AG (1998) Reinforcement learning: An introduction (MIT press).

6. Daw ND, Niv Y, & Dayan P (2005) Uncertainty-based competition between prefrontal and dorsolateral striatal systems for behavioral control. Nature neuroscience 8(12):1704-1711.

7. Schultz W, Dayan P, & Montague PR (1997) A neural substrate of prediction and reward. Science 275(5306):1593–1599.

8. Daw ND, O'Doherty JP, Dayan P, Seymour B, & Dolan RJ (2006) Cortical substrates for exploratory decisions in humans. Nature 441(7095):876–879.

9. Kemp JM & Powell T (1971) The connexions of the striatum and globus pallidus: synthesis and speculation. Philosophical Transactions of the Royal Society of London B: Biological Sciences 262(845):441–457.

10. Bogacz R & Gurney K (2007) The basal ganglia and cortex implement optimal decision making between alternative actions. Neural computation 19(2):442–477.

11. Wiecki TV & Frank MJ (2013) A computational model of inhibitory control in frontal cortex and basal ganglia. Psychological Review 120(2):329.

12. Hikosaka O, Kim HF, Yasuda M, & Yamamoto S (2014) Basal ganglia circuits for reward value-guided behavior. Annual review of neuroscience 37:289.

13. Frank MJ, et al. (2015) fMRI and EEG predictors of dynamic decision parameters during human reinforcement learning. The Journal of Neuroscience 35(2):485–494.

14. Ding L (2015) Distinct dynamics of ramping activity in the frontal cortex and caudate nucleus in monkeys. Journal of neurophysiology 114(3):1850–1861.

15. Haber SN (2003) The primate basal ganglia: parallel and integrative networks. Journal of chemical neuroanatomy 26(4):317–330.

16. Alexander GE, DeLong MR, & Strick PL (1986) Parallel organization of functionally segregated circuits linking basal ganglia and cortex. Annual review of neuroscience 9(1):357–381.

17. Haber SN, Kim K-S, Mailly P, & Calzavara R (2006) Reward-related cortical inputs define a large striatal region in primates that interface with associative cortical connections, providing a substrate for incentive-based learning. The Journal of Neuroscience 26(32):8368–8376.

18. Haber SN & Knutson B (2010) The reward circuit: linking primate anatomy and human imaging. Neuropsychopharmacology 35(1):4–26.

19. Damoiseaux J, et al. (2006) Consistent resting-state networks across healthy subjects. Proceedings of the national academy of sciences 103(37):13848–13853.

20. Bullmore E & Sporns O (2009) Complex brain networks: graph theoretical analysis of structural and functional systems. Nature Reviews Neuroscience 10(3):186–198.

21. Bressler SL & Menon V (2010) Large-scale brain networks in cognition: emerging methods and principles. Trends in cognitive sciences 14(6):277–290.

22. Power JD, et al. (2011) Functional network organization of the human brain. Neuron 72(4):665–678.

23. Medaglia JD, Lynall M-E, & Bassett DS (2015) Cognitive Network Neuroscience. Journal of cognitive neuroscience.

24. Liu X & Duyn JH (2013) Time-varying functional network information extracted from brief instances of spontaneous brain activity. Proceedings of the National Academy of Sciences 110(11):4392–4397.

25. Smith SM, et al. (2012) Temporally-independent functional modes of spontaneous brain activity. Proceedings of the National Academy of Sciences 109(8):3131–3136.

26. Allen EA, et al. (2012) Tracking whole-brain connectivity dynamics in the resting state. Cerebral cortex:bhs 352.

27. Kopell NJ, Gritton HJ, Whittington MA, & Kramer MA (2014) Beyond the connectome: the dynome. Neuron 83(6):1319–1328.

28. Mucha PJ, Richardson T, Macon K, Porter MA, & Onnela J-P (2010) Community structure in time-dependent, multiscale, and multiplex networks. Science 328(5980):876–878.

29. Bassett DS, et al. (2011) Dynamic reconfiguration of human brain networks during learning. Proceedings of the National Academy of Sciences 108(18):7641–7646.

30. Bassett DS, Yang M, Wymbs NF, & Grafton ST (2015) Learning-induced autonomy of sensorimotor systems. Nature Neuroscience.

31. Braun U, et al. (2015) Dynamic reconfiguration of frontal brain networks during executive cognition in humans. Proceedings of the National Academy of Sciences 112(37):11678–11683.

32. Bassett DS, et al. (2013) Task-based core-periphery organization of human brain dynamics. PLoS computational biology 9(9):e1003171.

33. Myers CE, et al. (2003) Dissociating medial temporal and basal ganglia memory systems with a latent learning task. Neuropsychologia 41(14):1919–1928.

34. Foerde K, Braun EK, & Shohamy D (2012) A trade-off between feedback-based learning and episodic memory for feedback events: evidence from Parkinson's disease. Neurodegenerative Diseases 11(2):93–101.

35. Foerde K & Shohamy D (2011) Feedback timing modulates brain systems for learning in humans. The Journal of Neuroscience 31(37):13157–13167.

36. Bates D, Mächler M, Bolker B, & Walker S (2015) Fitting Linear Mixed-Effects Models Usinglme4. Journal of Statistical Software 67(1).

37. Gershman SJ (2016) Empirical priors for reinforcement learning models. Journal of Mathematical Psychology 71:1–6.

38. Carpenter B, et al. (2015) Stan: a probabilistic programming language. Journal of Statistical Software.

39. Padoa-Schioppa C & Assad JA (2006) Neurons in the orbitofrontal cortex encode economic value. Nature 441(7090):223–226.

40. Braun U, et al. (2016) Dynamic brain network reconfiguration as a potential schizophrenia genetic risk mechanism modulated by NMDA receptor function. Proceedings of the National Academy of Sciences 113(44):12568–12573.

41. Kehoe EJ & Gormezano I (1980) Configuration and combination laws in conditioning with compound stimuli. Psychological Bulletin 87(2):351.

42. Wimmer GE, Daw ND, & Shohamy D (2012) Generalization of value in reinforcement learning by humans. European Journal of Neuroscience 35(7):1092–1104.

43. Wimmer GE & Shohamy D (2012) Preference by Association: How Memory Mechanisms in the Hippocampus Bias Decisions. Science 338(6104):270–273.

44. Eichenbaum H (2000) A cortical–hippocampal system for declarative memory. Nature Reviews Neuroscience 1(1):41–50.

45. Staresina BP & Davachi L (2009) Mind the gap: binding experiences across space and time in the human hippocampus. Neuron 63(2):267–276.

46. Palla K, Caron F, & Teh YW (2016) Bayesian nonparametrics for Sparse Dynamic Networks. arXiv preprint arXiv:1607.01624.

47. Durante D, Mukherjee N, & Steorts RC (2016) Bayesian Learning of Dynamic Multilayer Networks. arXiv preprint arXiv:1608.02209.

48. Shine JM, et al. (2016) The Dynamics of Functional Brain Networks: Integrated Network States during Cognitive Task Performance. Neuron 92(2):544–554.

49. Khambhati AN, Davis KA, Lucas TH, Litt B, & Bassett DS (2016) Virtual cortical resection reveals push-pull network control preceding seizure evolution. Neuron 91(5):1170–1182.

50. Siebenhühner F, Weiss SA, Coppola R, Weinberger DR, & Bassett DS (2013) Intra-and inter-frequency brain network structure in health and schizophrenia. PloS one 8(8):e72351.

51. Davidow JY, Foerde K, Galván A, & Shohamy D (2016) An Upside to Reward Sensitivity: The Hippocampus Supports Enhanced Reinforcement Learning in Adolescence. Neuron 92(1):93–99.

52. Foerde K, Race E, Verfaellie M, & Shohamy D (2013) A role for the medial temporal lobe in feedback-driven learning: evidence from amnesia. The Journal of Neuroscience 33(13):5698–5704.

53. Smith SM, et al. (2004) Advances in functional and structural MR image analysis and implementation as FSL. Neuroimage 23:S208-S219.

54. Bullmore ET & Bassett DS (2011) Brain graphs: graphical models of the human brain connectome. Annual review of clinical psychology 7:113–140.

55. Newman ME (2004) Fast algorithm for detecting community structure in networks. Physical review E 69(6):066133.

56. Bassett DS, Porter MA, Wymbs NF, Grafton ST, & Carlson JM (2013) Robust detection of dynamic community structure in networks. Chaos 23:013142.

57. Benjamini Y & Hochberg Y (1995) Controlling the false discovery rate: a practical and powerful approach to multiple testing. Journal of the Royal Statistical Society. Series B (Methodological):289-300.

58. Ishii S, Yoshida W, & Yoshimoto J (2002) Control of exploitation–exploration meta-parameter in reinforcement learning. Neural networks 15(4):665–687.

59. Andersson J, Smith S, & Jenkinson M (2008) Fnirt-fmrib's non-linear image registration tool. Human Brain Mapping 2008.

60. Satterthwaite TD, et al. (2012) Impact of in-scanner head motion on multiple measures of functional connectivity: relevance for studies of neurodevelopment in youth. Neuroimage.

61. Power JD, Barnes KA, Snyder AZ, Schlaggar BL, & Petersen SE (2012) Spurious but systematic correlations in functional connectivity MRI networks arise from subject motion. Neuroimage 59(3):2142–2154.

62. Satterthwaite TD, et al. (2012) An Improved Framework for Confound Regression and Filtering for Control of Motion Artifact in the Preprocessing of Resting-State Functional Connectivity Data. NeuroImage.

63. Holme P & Saramäki J (2011) Temporal Networks. Physics Reports 519(3):97–125.

64. Kivelä M, et al. (2014) Multilayer networks. Journal of Complex Networks 2(3):203–271.

65. Porter MA, Onnela J-P, & Mucha PJ (2009) Communities in networks. Notices of the AMS 56(9):1082–1097.

66. Fortunato S (2010) Community detection in graphs. Physics reports 486(3):75–174.

67. Meunier D, Lambiotte R, & Bullmore ET (2010) Modular and hierarchically modular organization of brain networks. Frontiers in neuroscience 4:200.

68. Cole MW, Bassett DS, Power JD, Braver TS, & Petersen SE (2014) Intrinsic and task-evoked network architectures of the human brain. Neuron 83(1):238–251.

69. Yeo BT, et al. (2011) The organization of the human cerebral cortex estimated by intrinsic functional connectivity. Journal of neurophysiology 106(3):1125–1165.

70. Jutla IS, Jeub LG, & Mucha PJ (2011) A generalized Louvain method for community detection implemented in MATLAB. URL http://netwiki.amath.unc.edu/GenLouvain.

71. Good BH, de Montjoye Y-A, & Clauset A (2010) Performance of modularity maximization in practical contexts. Physical Review E 81(4):046106.

72. Betzel RF, Satterthwaite TD, Gold JI, & Bassett DS (2016) A positive mood, a flexible brain. arXiv preprint arXiv:1601.07881.

73. Gelman A (2006) Prior distributions for variance parameters in hierarchical models (comment on article by Browne and Draper). Bayesian analysis 1(3):515–534.

74. Lewandowski D, Kurowicka D, & Joe H (2009) Generating random correlation matrices based on vines and extended onion method. Journal of multivariate analysis 100(9):1989–2001.

